# A free-living protist that lacks canonical eukaryotic DNA replication and segregation systems

**DOI:** 10.1101/2021.03.14.435266

**Authors:** Dayana E. Salas-Leiva, Eelco C. Tromer, Bruce A. Curtis, Jon Jerlström-Hultqvist, Martin Kolisko, Zhenzhen Yi, Joan S. Salas-Leiva, Lucie Gallot-Lavallée, Geert J. P. L. Kops, John M. Archibald, Alastair G. B. Simpson, Andrew J. Roger

## Abstract

Cells must replicate and segregate their DNA with precision. In eukaryotes, these processes are part of a regulated cell-cycle that begins at S-phase with the replication of DNA and ends after M-phase. Previous studies showed that these processes were present in the last eukaryotic common ancestor and the core parts of their molecular systems are conserved across eukaryotic diversity. However, some unicellular parasites, such as the metamonad *Giardia intestinalis*, have secondarily lost components of the DNA processing and segregation apparatuses. To clarify the evolutionary history of these systems in these unusual eukaryotes, we generated a high-quality draft genome assembly for the free-living metamonad *Carpediemonas membranifera* and carried out a comparative genomics analysis. We found that parasitic and free-living metamonads harbor a conspicuously incomplete set of canonical proteins for processing and segregating DNA. Unexpectedly, *Carpediemonas* species are further streamlined, lacking the origin recognition complex, Cdc6 and other replisome components, most structural kinetochore subunits including the Ndc80 complex, as well as several canonical cell-cycle checkpoint proteins. *Carpediemonas* is the first eukaryote known to have lost this large suite of conserved complexes, suggesting that it has a highly unusual cell cycle and that unlike any other known eukaryote, it must rely on novel or alternative set of mechanisms to carry out these fundamental processes.

---

DNA replication, repair and segregation are critically important and conserved processes in eukaryotes that have been intensively studied in model organisms^1^. The initial step of DNA replication is accomplished by the replisome, a set of highly conserved proteins that is tightly regulated to minimize mutations^2^. The replisome relies on the interactions between cis-acting DNA sequences and trans-acting factors that serve to separate the template and promote RNA-primed DNA synthesis. This occurs by the orderly assembly of the origin recognition (ORC), the pre-replicative (pre-RC), pre-initiation (pre-IC) and replication progression (RPC) complexes^3–6^. The synthesis of DNA usually encounters disruptive obstacles as replication proceeds and can be rescued either through template switching via trans-lesion or recombination-dependent synthesis. Trans-lesion synthesis uses replicative and non-replicative DNA polymerases to by-pass the lesion through multiple strategies that incorporate nucleotides opposite to it^7^, while recombination-dependent synthesis uses non-homologous or homologous templates for repair (reviewed in refs.^8, 9^). Recombination-dependent synthesis occurs in response to single- or double-strand DNA breakage^8, 10, 11^. Other repair mechanisms occur throughout the cell cycle, fixing single-strand issues through base excision, nucleotide excision or mismatch repair, but they may also be employed during replication depending on the source of the damage. All of the repair processes are overseen by multiple regulation checkpoints that permit or stall DNA replication and the progression of the cell cycle. During M-phase the replicated DNA has to form attachments with the microtubule-based spindle apparatus via kinetochores, large multi-subunit complexes built upon centromeric chromatin^12^. Unattached kinetochores catalyse the formation of a soluble inhibitor of the cell cycle, preventing precocious chromosome segregation, a phenomenon known as the spindle assembly checkpoint (SAC)^12^. Failure to pass any of these checkpoints (*e.g.*, G1/S, S, G2/M and SAC checkpoints reviewed in refs.^12–14^) leads to genome instability and may result in cell death.

To investigate the diversity of DNA replication, repair, and segregation processes, we conducted a eukaryote-wide comparative genomics analysis with a special focus on metamonads, a major protist lineage comprised of parasitic and free-living anaerobes. Parasitic metamonads such as *Giardia intestinalis* and *Trichomonas vaginalis* are extremely divergent from model system eukaryotes, exhibit a diversity of cell division mechanisms (*e.g.*, closed/semi-open mitosis), possess metabolically reduced mitosomes or hydrogenosomes instead of mitochondria, and lack several canonical eukaryotic features on the molecular and genomic-level^15–17^. Indeed, recent studies show that metamonad parasites have secondarily lost parts of the ancestral DNA replication and segregation apparatuses^18, 19^. Furthermore, metamonad proteins are often highly divergent compared to other eukaryotic orthologs, indicating a high substitution rate in these organisms that is suggestive of error-prone replication and/or DNA repair^20, 21^. Yet, it is unclear whether the divergent nature of proteins studied in metamonads is the result from the host-associated lifestyle or is a more ancient feature of Metamonada. To increase the representation of free-living metamonads in our analyses, we have generated a high-quality draft genome assembly of *Carpediemonas membranifera, a* flagellate isolated from hypoxic marine sediments^22^. Our analyses of genomes from across the tree of eukaryotes show that many systems for DNA replication, repair, segregation, and cell cycle control are ancestral to eukaryotes and highly conserved. However, metamonads have secondarily lost an extraordinarily large number of components. Most remarkably, the free-living *Carpediemonas* species have been drastically reduced further, having lost a large set of key proteins from the replisome and cell-cycle checkpoints (*i.e.*, including several from the kinetochore and repair pathways). We propose a hypothesis of how DNA replication may be achieved in this organism.

## Results

### The *C. membranifera* genome assembly is complete

Our assembly for *C. membranifera* (a member of the Fornicata clade within metamonads, **Fig. 1**) is highly contiguous (**Table 1**) and has deep read coverage (*i.e.*, median coverage of 150× with short reads and 83× with long-reads), with an estimated genome completeness of 99.27% based on the Merqury^23^ method. 97.6% of transcripts mapped to the genome along their full length with an identity of ≥ 95% while a further 2.04% mapped with an identity between 90 - 95%. The high contiguity of the assembly is underscored by the large number of transcripts mapped to single contigs (90.2%), and since the proteins encoded by transcripts were consistently found in the predicted proteome, the latter is also considered to be of high quality. We also conducted BUSCO analyses, with the foreknowledge that genomic streamlining typical in Metamonada has led to the loss of many conserved proteins^16, 17, 24^. Our analyses show that previously completed metamonad genomes only encoded between 60% to 91% BUSCO proteins, while *C. membranifera* exhibits a relatively high 89% (**Table 1**, **Supplementary Information**). In any case, our coverage estimates for the *C. membranifera* genome for short and long read sequencing technologies are substantially greater than those found to be sufficient to capture genic regions that otherwise would had been missed (*i.e.*, coverage >52× for long reads and >60× for short paired-end reads, see ref.^25^). All these various data indicate that the draft genome of *C. membranifera* is nearly complete; if any genomic regions are missing, they are likely confined to difficult-to-sequence highly repetitive regions such as telomeres and centromeres.

**Figure 1.**
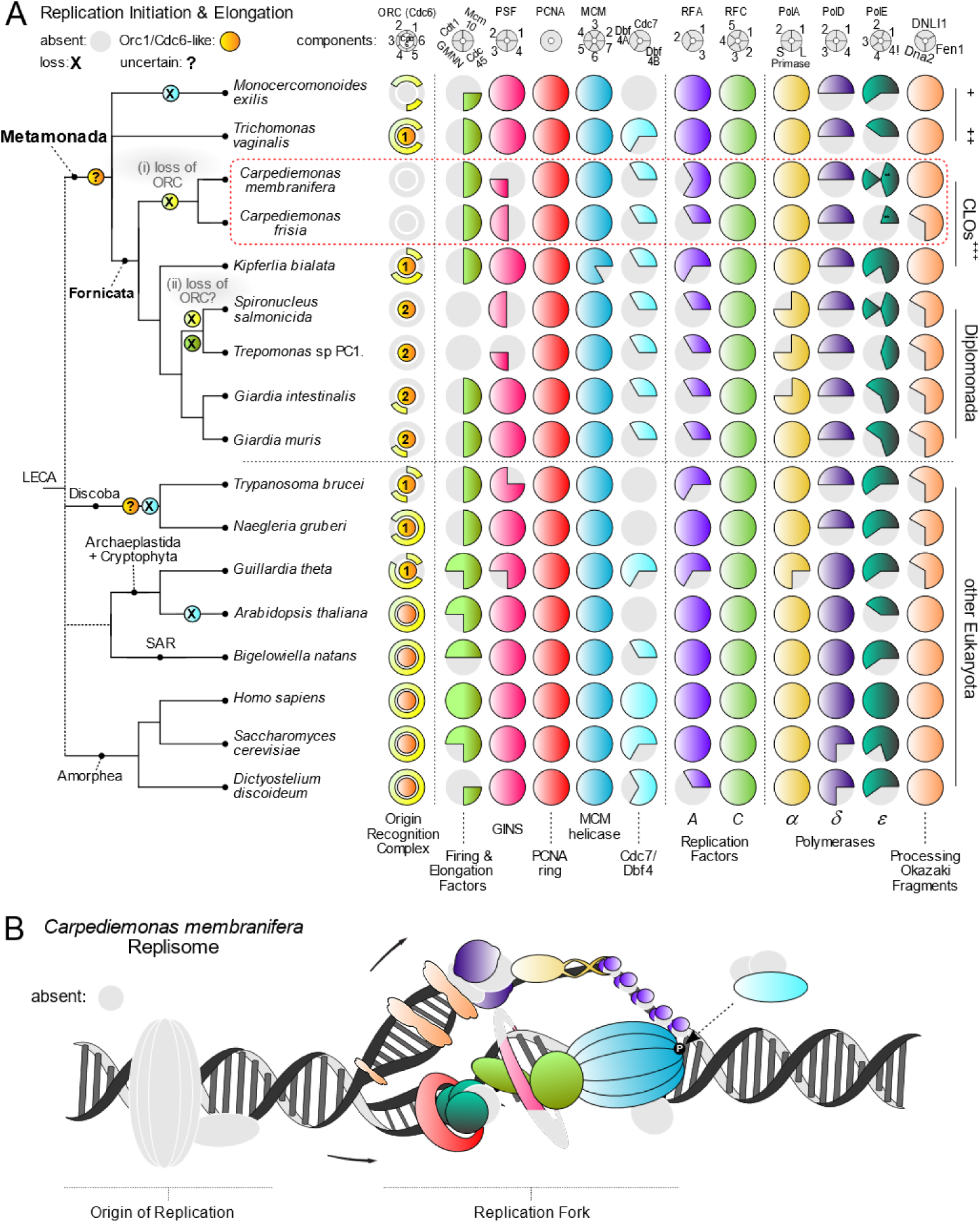
The distribution of core molecular systems in the replisome and DNA repair across eukaryotic diversity. A schematic global eukaryote phylogeny is shown on the left with classification of the major metamonad lineages indicated at right. **A)** The Replisome. Reduction of the replication machinery complexity and extensive loss of the Orc1-6 subunits are observed in metamonad lineages, including the unexpected loss of the highly conserved ORC complex and Cdc6 in *Carpediemonas*. Most metamonad Orc1 and Cdc6 homologs were conservatively named as ‘Orc1/Cdc6-like’ as they are very divergent, do not have the typical domain architecture and, in phylogenetic reconstructions, they form clades separate from the main eukaryotic groups, preventing confident orthology assignments (**Supplementary Figure 1**). Numbers within subunits represent the number of copies and are only presented for ORC components, additional information in **Supplementary Table 1**. The polymerase epsilon (ε) is composed of 4 subunits, but we included the interacting protein Chrac1 (depicted as ‘4!’ in the figure) as its HMM retrieves the polymerase delta subunit Dbp3 from *S. cerevisiae*. *Firing and elongation factors, **Protein fusion between the catalytic subunit and subunit 2 of DNA polymerase ε. ^+^ Preaxostyla, ^++^ Parabasalida, **^+++^***Carpediemonas*-Like Organisms. **B)** Predicted *Carpediemonas* replisome overlayed on a typical eukaryotic replisome. Origin recognition (ORC), Cdc6 and replication progression (RPC) complexes are depicted. Grey colour represents the absence of typical eukaryotic proteins in *C. membranifera* replisome.

**Table 1.**
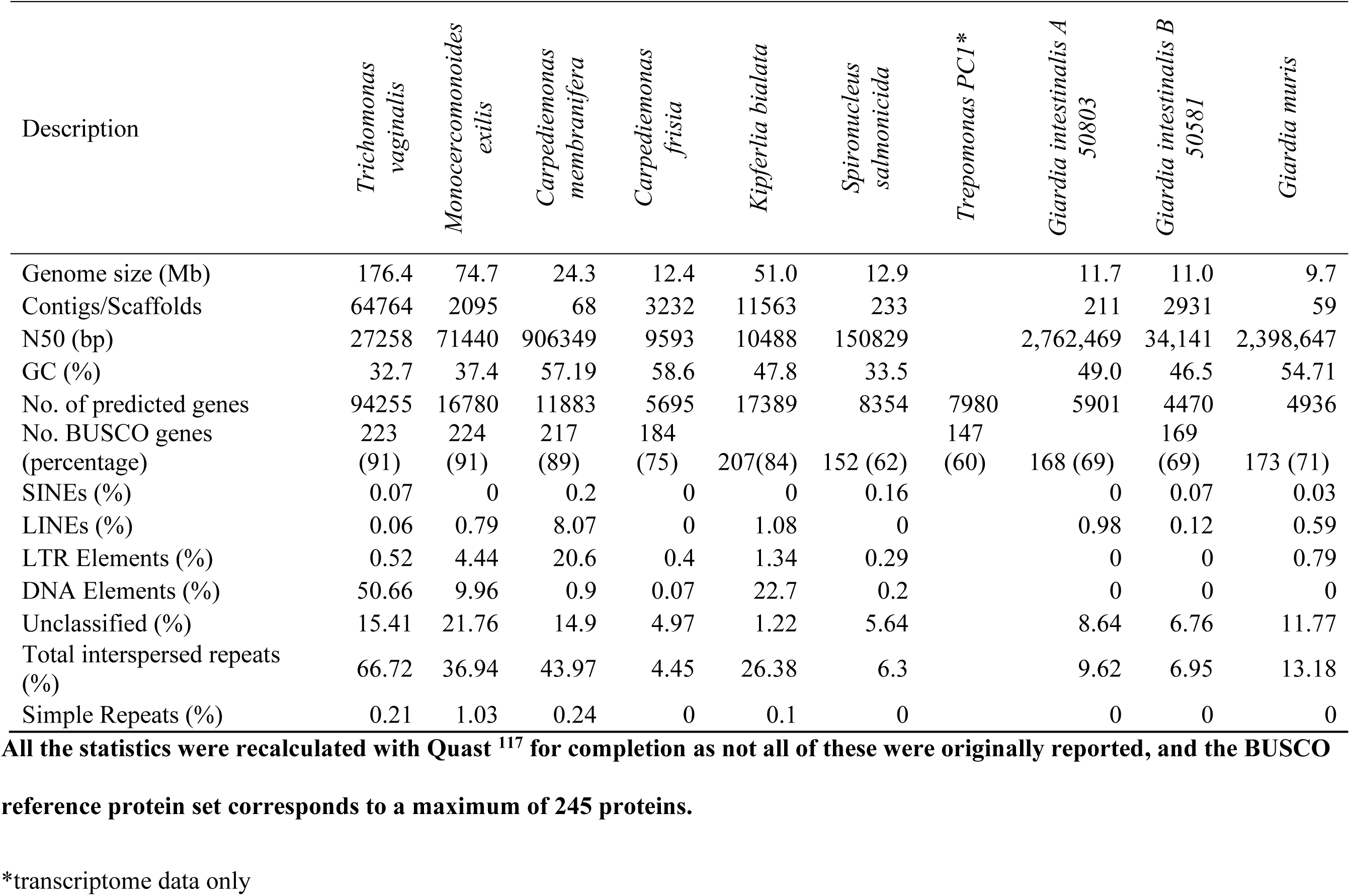
Summary statistics of nuclear genomes of Metamonada species.

Note that a previous study conducted a metagenomic assembly of a related species, *Carpediemonas frisia,* together with its associated prokaryotic microbiota^26^. For completeness, we have included these data in our comparative genomic analyses (**Table 1**, **Supplementary Information**), although we note that the *C. frisia* metagenomic bin is based on only short-read data and might be partial.

### Extreme streamlining of the DNA replication apparatus in metamonads

The first step in the replication of DNA is the assembly of ORC which serves to nucleate the pre-RC formation. The initiator protein Orc1first binds an origin of replication, followed by the recruitment of Orc 2-6 proteins, which associate with chromatin^27^. As the cell transitions to G1 phase, the initiator Cdc6 binds to the ORC, forming a checkpoint control^28^. Cdt1 then joins Cdc6, promoting the loading of the replicative helicase MCM forming the pre-RC, a complex that remains inactive until the onset of S-phase when the ‘firing’ factors are recruited to convert the pre-RC into the pre-IC^3–5^. Additional factors join to form the RPC to stimulate replication elongation^29^. While the precise replisome protein complement varies somewhat between different eukaryotes, metamonads show dramatic variation in ORC, pre-RC and replicative polymerases (**Fig. 1**). The presence-absence of ORC and Cdc6 proteins is notably patchy across Metamonada. Strikingly, whereas all most metamonads retain up to two paralogs of the core protein family Orc1/Cdc6 (here called Orc1 and Orc1/Cdc6-like, **Supplementary Figure 1**), plus some orthologs of Orc 2-6, all these proteins are absent in *C. membranifera* and *C. frisia* (**Fig. 1**, **Supplementary Table 1**). The lack of these proteins in a eukaryote is unexpected and unprecedented, since their absence would be expected to make the genome prone to DSBs and impair DNA replication, as well as interfere with other non-replicative processes^30^. To rule out false negatives, we conducted further analyses using metamonad-specific HMMs (Hidden Markov Models), various other profile-based search strategies (**Supplementary Information**), tBLASTn^31^ searches (*i.e.*, on the genome assembly and unassembled long-reads), and applied HMMER^32^ on 6-frame assembly translations. These additional methods were sufficiently sensitive to identify these proteins in all nuclear genomes we examined, with the exception of the *Carpediemonas* species and the highly reduced, endosymbiotically-derived nucleomorphs of cryptophytes and chlorarachniophytes (**Supplementary Information, Supplementary Table 1**, **Supplementary Fig. 1 and 2**). *Carpediemonas* species are, therefore, the only known eukaryotes to completely lack ORC and Cdc6.

**Figure 2.**
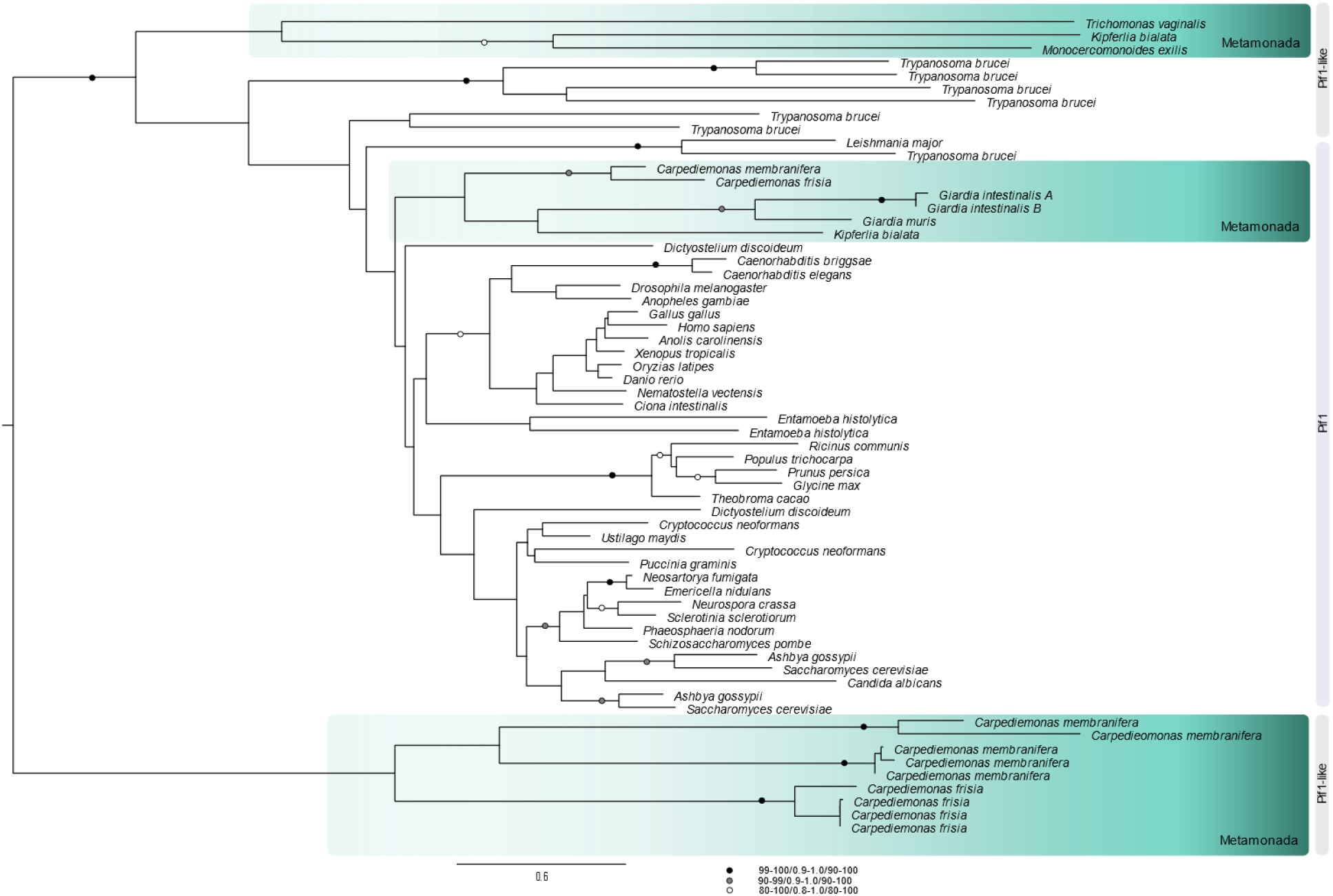
Pif1 protein family expansion. Pif1 helicase family tree. Three clades are highlighted: at the top, a Pif1-like clade encompassing some metamonads and at the bottom a *Carpediemonas-*specific Pif1-like clade. The third clade shows the typical Pif1 orthologs encompassing fornicates. The maximum-likelihood tree was inferred under the LG+PMSF(C60)+F+ Γ model using 100 bootstraps based on an alignment length of 265 sites. The tree was midpoint-rooted and the support values on the branches correspond to SH-aLRT/aBayes/standard bootstrap (values below 80/0.8/80 are not shown). The scale bar shows the inferred number of amino acid substitutions per site.

### DNA damage repair systems have undergone several modifications

DNA repair occurs continuously during the cell cycle depending on the type or specificity of the lesion. Among the currently known mechanisms are base-excision repair (BER), nucleotide excision repair (NER), mismatch repair (MMR), and double strand break repair, with the latter conducted by either homologous recombination (HR), canonical non-homologous end joining (NHEJ) or alternative end joining (a-EJ)^8, 14^. MMR can be coupled directly to replication or play a role in HR. MMR, BER and NER are present in all studied taxa (**Supplementary Table 1**), although our analyses indicate that damage sensing and downstream functions in NER seem to be modified in the metamonad taxa Parabasalia and Fornicata due to the absence of the XPG and XPC sensor proteins.

Double strand breaks (DSBs) are extremely dangerous for cells and can occur as a result of damaging agents or from self-inflicted cuts during DNA repair and meiosis. NHEJ requires the heterodimer Ku70-Ku80 to recruit the catalytic kinase DNA-PKcs and accessory proteins. Metamonads lack all of these proteins, as do a number of other eukaryotes investigated here and in ref.^33^. The a-EJ system seems to be fully present in metamonads like *C. membranifera*, partial in others, and completely absent in parasitic diplomonads. NHEJ is thought to be the predominant mechanism for repairing DSBs in eukaryotes^34^, but since our analyses indicate this pathway is absent in metamonads and a-EJ is highly mutagenic^8^, the HR pathway is likely to be essential for DSB repair in most metamonads. Repair by the HR system occurs through multiple sub-pathways that are influenced by the extent of the similarity of the DNA template or its flanking sequences to the sequences near the break. HR complexes are recruited during DNA replication and transcription, and utilize DNA, transcript-RNA or newly synthetized transcript-cDNA as a homologous template^11, 35–40^. These complexes are formed by recombinases from the RecA/Rad51 family that interact with members of the Rad52 family and chromatin remodeling factors of the SNF2/SWI2 sub-family^41, 42^. Although the recombinases Rad51A-D are all present in most eukaryotes, we found a patchy distribution in metamonads (**Supplementary Table 1**, **Supplementary Fig. 3**). All examined Fornicata have lost the major recombinase Rad51A and have two paralogs of the meiosis-specific recombinase Dmc1, as first noted in *Giardia intestinalis*^43^. Dmc1 has been reported to provide high stability to recombination due to strong D-loop resistance to strand dissociation^44^. The recombination mediator Rad52 is present in metamonads but Rad59 or Rad54 are not. Metamonads have no components of an ISWI remodeling complex yet retain a reduced INO80 complex. Therefore, replication fork progression and HR are likely to occur under the assistance of INO80 alone. HR requires endonucleases and exonucleases, and our searches for proteins additional to those from the MMR pathway revealed a gene expansion of the Flap proteins from the Rad2/XPG family in some metamonads. We also found proteins of the PIF1 helicase family that encompasses homologs that resolve R-loop structures, unwind DNA–RNA hybrids and assists in fork progression in regular replication and HR^45, 46^. Phylogenetic analysis reveals that although *Carpediemonas* species have orthologs that branch within a metamonad group in the main PIF1 clade (**Fig. 2**), they also possess a highly divergent clade of PIF1-like proteins. Each *Carpediemonas* species has multiple copies of PIF1-like proteins that have independently duplicated within each species; these may point to the *de novo* emergence of specialized functions in HR and DNA replication for these proteins. Metamonads appear capable of using all of the HR sub-pathways (*e.g.*, classical DSB repair, single strand annealing, break induced replication), but these are modified (**Supplementary Table 1**, **Supplementary Figure 3**). Overall, the presence-absence patterns of the orthologs involved in DSB repair in Fornicata point to the existence of a highly specialized HR pathway which is presumably not only essential for the cell cycle of metamonads but is also likely the major pathway for replication-related DNA repair and recombination.

### Modified DSB damage response checkpoints in metamonads

Checkpoints constitute a cascade of signaling events that delay replication until DNA lesions are resolved^13^. The ATR-Chk1, ATM-Chk2 and DNA-PKcs pathways are activated by the interaction of TopBP1 and the 9-1-1 complex (Rad9-Hus1-Rad1) for DNA repair regulation during replication stress and response to DSBs^47^. The ATR-Chk1 signaling pathway is the initial response to ssDNA damage and is responsible for the coupling of DNA replication with mitosis, but when it is defective, the ssDNA is converted into DSBs to activate the ATM-Chk2 pathway. The DNA-PKcs act as sensors of DSBs to promote NHEJ, but we found no homologs of DNA-PKcs in metamonads (**Supplementary Fig. 3**), which is consistent with the lack of a NHEJ repair pathway in the group. All the checkpoint pathways described are present in humans and yeasts, while the distribution of core checkpoint proteins in the remaining taxa is patchy. Notably, Fornicata lack several of the proteins thought to be needed to activate the signaling kinase cascades and, while orthologs of ATM or ATR kinases are present in some fornicates, there are no clear orthologs of Chk1 or Chk2 in metamonads except in *Monocercomonoides exilis* (**Supplementary Table 1**, **Supplementary Fig. 3**). *Carpediemonas* species and *Kipferlia bialata* contain ATM and ATR but lack Chk1, Chk2 and Rad9. Diplomonads possess none of these proteins. The depletion of Chk1 has been shown to increase the incidence of chromosomal breaks and mis-segregation^48^. All these absences reinforce the idea that the checkpoint controls in Fornicata are non-canonical.

### Reduction of mitosis and meiosis machinery in metamonads

Eukaryotes synchronize cell cycle progression with chromosome segregation by a kinetochore based signaling system called the spindle assembly checkpoint (SAC)^49, 50^ that is ancestral to all eukaryotes (**Fig. 3A, B**). Kinetochores primarily form microtubule attachments through the Ndc80 complex, which is connected through a large network of structural subunits to a histone H3-variant CenpA that is specifically deposited at centromeres^12^. To prevent premature chromosome segregation, unattached kinetochores catalyse the production of the Mitotic Checkpoint Complex (MCC)^49^, a cytosolic inhibitor of the Anaphase Promoting Complex/Cyclosome (APC/C), a large multi-subunit E3 ubiquitin ligase that drives progression into anaphase by promoting the proteolysis of its substrates such as various Cyclins^51^ (**Fig. 3A**). Our analysis indicates the reduction of ancestral complexity of these proteins in metamonads (**Fig. 3C**, **Supplementary Table 1**, **Supplementary Fig. 4**). Surprisingly, such reduction is most extensive in *Carpediemonas* species. We found that most structural kinetochore subunits, a microtubule plus-end tracking complex and all four subunits of the Ndc80 complex are absent (**Fig. 3C**, **Supplementary Fig. 4**). None of our additional search strategies led to the identification of Ndc80 complex members, making *Carpediemonas* the only known eukaryotic lineage without it, except for kinetoplastids, which appear to have lost the canonical kinetochore and replaced it by an analogous molecular system, although there is still some controversy about this loss^52, 53^. With such widespread absence of kinetochore components it might be possible that *Carpediemonas* underwent a similar replacement process to that of kinetoplastids^52^. We did however find a potential candidate for the centromeric Histone H3-variant (CenpA) in *C. membranifera*. CenpA forms the basis of the canonical kinetochore in most eukaryotes^54^ (**Supplementary Fig. 5**). On the other hand, the presence or absence of CenpA is often correlated with the presence/absence of its direct interactor CenpC^19^. Similar to diplomonads, *C. membranifera* lacks CenpC and therefore the molecular network associated with kinetochore assembly on CenpA chromatin may be very different.

**Figure 3.**
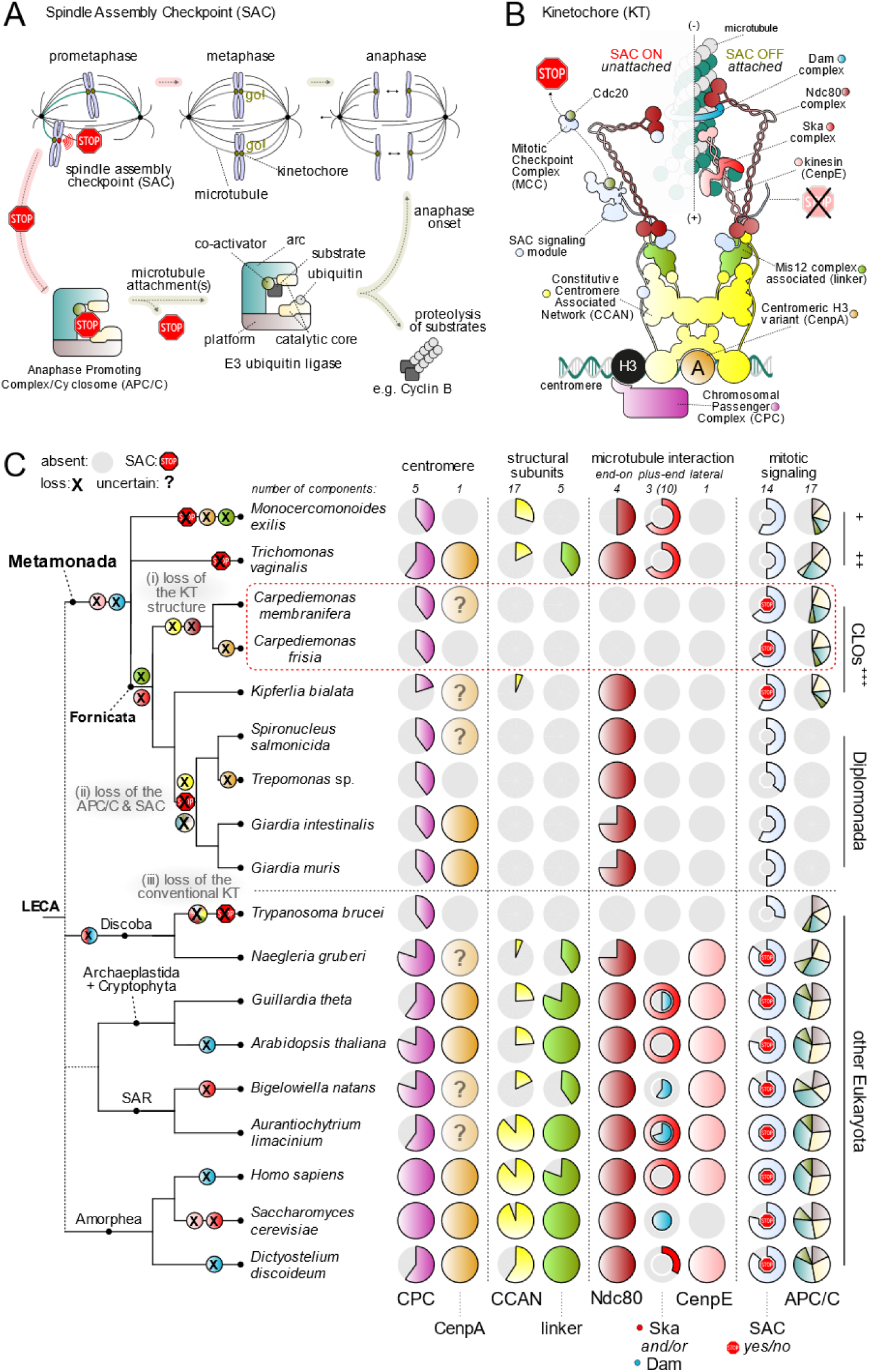
Radical reduction of ancestral kinetochore network complexity in *Carpediemonas* species. **A)** Schematic of canonical mitotic cell cycle progression in eukaryotes. During mitosis, duplicated chromosomes each attach to microtubules (MTs) emanating from opposite poles of the spindle apparatus, in order to be segregated into two daughter cells. Kinetochores (KTs) are built upon centromeric DNA to attach microtubules to chromosomes. To prevent precocious chromosome segregation, unattached KTs signal to halt cell cycle progression (STOP), a phenomenon known as the Spindle Assembly Checkpoint (SAC). The SAC entails the inhibition of the Anaphase Promoting Complex/Cyclosome (APC/C), a multi-subunit E3 ubiquitin ligase complex that drives the entry of mitotic cells into anaphase by promoting the proteolysis of its substrates. Once all KTs are correctly attached to spindle MTs and aligned in the middle of the cell (metaphase), the APC/C is released, its substrates are degraded, and chromosome segregation is initiated (anaphase). **B)** Cartoon of the molecular makeup of a single KT unit that was likely present in Last Eukaryotic Common Ancestor (LECA). Colours indicate the various functional complexes and structures. The primary KT structure is provided by the Constitutive Centromere Associated Network (CCAN; yellow), which is built upon centromeric chromatin that contains Centromere protein A (CenpA; orange), a centromere-specific Histone H3. During mitosis the CCAN recruits the Mis12 complex (linker; light green), which provides a platform for the recruitment of the SAC signalling (light blue) and microtubule-interacting complexes. The Chromosomal Passenger Complex (CPC; dark purple) localizes at the inner centromere and harbours a kinase (aurora) that regulates microtubule attachments. Unattached KTs catalyse the production of a diffusible cytosolic inhibitor of the APC/C, known as the mitotic checkpoint complex (MCC), which captures the mitotic APC/C co-activator Cdc20. Initial KT-MT encounters are driven by the kinesin Centromere protein E (CenpE; pink), which binds MTs at the lateral sides. The Ndc80 complex (dark red) constitutes the main end-on MT binding activity of KTs. To facilitate the tracking of the plus-end (+) of MT during anaphase, eukaryotes utilize two different complexes: Dam (light purple; likely not present in LECA) and Ska (red). Once KTs are bound by MTs, SAC signalling proteins are removed and the SAC is turned off. **C)** Reconstruction of the evolution of the KT and mitotic signalling in eukaryotes based on protein presence-absence patterns reveals extensive reduction of ancestral KT complexity and loss of the SAC in most metamonad lineages, including the striking loss of the highly conserved core MT-binding activity of the KT (Ndc80) in *Carpediemonas*. On top/bottom of panel C: the number of components per complex and different structural parts of the KT, SAC signalling and the APC/C. Middle: presence/absence matrix of KT, SAC and APC/C complexes; one circle per complex, colours correspond to panel A & B; grey indicates its (partial) loss (for a complete overview see **Supplementary Table 1**, **Supplementary Fig. 4)**. The red STOP sign indicates the likely presence of a functional SAC response (see for discussion **Supplementary** Fig. 6). On the left: cartoon of a phylogenetic tree of metamonad and other selected eukaryotic species with a projection of the loss and gain events on each branch. Specific loss events of kinetochore and SAC genes in specific lineages are highlighted in colour.

Most metamonads encode all MCC components, but diplomonads lost the SAC response and the full APC/C complex^55^. In contrast, only *Carpediemonas* species and *K. bialata* have MCC subunits that contain the conserved short linear motifs to potentially elicit a canonical SAC signal^51, 56^ (**Supplementary Fig. 6**). Interestingly, not all of these motifs are present, and most are seemingly degenerate compared to their counterparts in other eukaryotic lineages (**Supplementary Fig. 6C**). Also, many other SAC-related genes are conserved, even in diplomonads (*e.g.*, Mad2, MadBub)^55^. Furthermore, the cyclins in *C. membranifera*, the main target of SAC signalling, have a diverged destruction motif (D-box) in their N-termini (**Supplementary Fig. 6C**). Collectively, our observations indicate that *Carpediemonas* species could elicit a functional SAC response, but whether this would be kinetochore-based is unclear. Alternatively, SAC-related genes could have been repurposed for another cellular function(s) as in diplomonads^55^. Given that ORC has been observed to interact with the kinetochore (throughout chromosome condensation and segregation), centrioles and promotes cytokinesis^30^, the lack of Ncd80 and ORC complexes suggest that *Carpediemonas* species possess radically unconventional cell division systems.

Neither sexual nor parasexual processes have been directly observed in Metamonada^43^. Nonetheless, our surveys confirm the conservation of the key meiotic proteins in metamonads^43^, including Hap2 (for plasmogamy) and Gex1 (karyogamy). Unexpectedly, *Carpediemonas* species have homologs from the tmcB family that acts in the cAMP signaling pathway specific for sexual development in *Dictyostelium*^57^, and sperm-specific channel subunits (*i.e.*, CatSper α, β, δ and γ) reported previously only in Opisthokonta and three other protists^58^. In opisthokonts, the CatSper subunits enable the assembly of specialized Ca^2+^ influx channels and are involved in the signaling for sperm maturation and motility^58^. In *Carpediemonas,* the tmcB family and CatSper subunits could similarly have a role in signaling and locomotion pathways required for a sexual cycle. As proteins in the cAMP pathway and Ca^2+^ signaling cooperate to generate a variety of complex responses, the presence of these systems in *Carpediemonas* species but absence in all other sampled metamonads is intriguing and deserves further investigation. Even if these systems are not directly involved in a sexual cycle, the presence of Hap2 and Gex1 proteins is strong evidence that *C. membranifera* can reproduce sexually. Interestingly, based on the frequencies of single nucleotide polymorphisms, *C. membranifera* is predicted to be haploid (**Supplementary Fig. 7**). If this is correct, its sexual reproduction should include the formation of a zygote followed by a meiotic division to regain its haploid state^59^.

### Acquisition of DNA replication and repair proteins in *Carpediemonas* by lateral gene transfer

The unprecedented absence of many components of canonical DNA replication, repair, and segregation systems in *Carpediemonas* species led us to investigate whether they had been replaced by analogous systems acquired by lateral gene transfer (LGT) from viruses or prokaryotes. We detected four Geminivirus-like replication initiation protein sequences in the *C. membranifera* genome but not in *C. frisia*, and helitron-related helicase endonucleases in both *Carpediemonas* genomes. All these genes were embedded in high-coverage eukaryotic scaffolds, yet all of them lack introns and show no evidence of gene expression in the RNA-Seq data. As RNA was harvested from log-phase actively replicating cell cultures, their lack of expression suggests it is unlikely that these acquired proteins were coopted to function in the replication of the *Carpediemonas* genomes.

Nevertheless, the presence of Geminivirus protein-coding genes is intriguing as these viruses are known, in other systems (*e.g.*, plants, insects), to alter host transcriptional controls and reprogram the cell-cycle to induce the host DNA replication machinery^60, 61^. We also detected putative LGTs of Endonuclease IV, RarA and RNAse H1 from prokaryotes into a *Carpediemonas* ancestor (**Supplementary Information, Supplementary Fig. 8, 9 and 10**). Of these, RarA is ubiquitous in bacteria and eukaryotes and acts during replication and recombination in the context of collapsed replication forks^62, 63^. Interestingly, *Carpediemonas* appears to have lost the eukaryotic ortholog, and only retains the acquired prokaryotic-like RarA, a gene that is expressed (*i.e.*, transcripts are present in the RNA-Seq data). RNAse Hs are involved in the cleavage of RNA from RNA:DNA hybrid structures that form during replication, transcription, and repair, and, while eukaryotes have a monomeric RNAse H1 and a heterotrimeric RNAse H2, prokaryotes have either one or both types.

Eukaryotic RNAse H1 removes RNA primers during replication and R-loops during transcription, and also participates in HR-mediated DSB repair^64, 65^. The prokaryotic homologs have similar roles during replication and transcription^66^. *C. membranifera* lacks a typical eukaryotic RNAse H1 but has two copies of prokaryotic homologs. Both are located in scaffolds comprising intron-containing genes and have RNA-Seq coverage, clearly demonstrating that they are not from prokaryotic contaminants in the assembly.

## Discussion

### Genome streamlining in metamonads

The reductive evolution of the DNA replication and repair, and segregation systems and the low retention of proteins in the BUSCO dataset in metamonads demonstrate that substantial gene loss has occurred (**Supplementary information**), providing additional evidence for streamlining of gene content prior to the last common ancestor of Metamonada^15–17^. However, the patchy distribution of genes within the group suggests ongoing differential reduction in different metamonad groups. Such reduction – especially the unprecedented complete absence of systems such as the ORC, Cdc6 and kinetochore Ndc80 complexes in *Carpediemonas* species – demands an explanation. Whereas the loss of genes from varied metabolic pathways is well known in lineages with different lifestyles^67–72^, loss of cell cycle, DNA damage sensing and repair genes in eukaryotes is extremely rare. New evidence from yeasts of the genus *Hanseniaspora* suggests that the loss of proteins in these systems can lead to genome instability and long-term hypermutation leading to high rates of sequence substitution^67^. This could also apply to metamonads, especially fornicates, which are well known to have undergone rapid sequence evolution; these taxa form a highly divergent clade with very long branches in phylogenetic trees^20, 73^. Most of the genes that were retained by Metamonada in the various pathways we examined were divergent in sequence relative to homologs in other eukaryotes and many of the gene losses correspond to proteins that are essential in model system eukaryotes. Gene essentiality appears to be relative and context-dependent, and some studies have shown that the loss of ‘indispensable’ genes could be permitted by evolving divergent pathways that provide similar activities via chromosome stoichiometry changes and compensatory gene loss^67–69, 74^.

The patchy distribution of genes from different ancestral eukaryotic pathways suggests that the last common ancestor of Metamonada had a broad gene repertoire for maintaining varied metabolic functions under fluctuating environmental conditions offered by diverse oxygen-depleted habitats. Although the loss of proteins and genomic streamlining are well known in parasitic diplomonads^15, 16^, the Fornicata, as a whole, tend to have a reduced subset of the genes that are commonly found in core eukaryotic pathways. In general, such gene content reduction can partially be explained as the result of historical and niche-specific adaptations^75^. Yet, given that 1) genome maintenance mostly depends on the cell cycle checkpoints, DNA repair pathways, and their interactions^14, 76^, 2) the lack of several proteins related to these pathways that were present in the last common ancestor of metamonads, 3) aneuploidy and high overall rates of sequence evolution have been observed in metamonads ^77, 78^, and, 4) the loss of DNA repair genes can be associated with substantial gene loss and sequence instability that apparently boosts the rates of sequence evolution^67^, it is likely that genome evolution in the Fornicata clade has been heavily influenced by their error-prone DNA maintenance mechanisms.

### Non-canonical replication initiation and replication licensing in *Carpediemonas*

Origin-independent replication has been observed in the context of DNA repair (reviewed in ref.^10^) and in origin-deficient or -depleted chromosomes in yeast^79^. These studies have highlighted the lack of (or reduction in) the recruitment of ORC and Cdc6 onto the DNA, but no study to date has documented regular eukaryotic DNA replication in the absence of genes encoding these proteins. While it is possible that extremely divergent versions of ORC and Cdc6 are governing the recognition of origins of replication and replication licensing in *Carpediemonas* species, we have no evidence for this. Instead, our findings suggest the existence of an as-yet undiscovered underlying eukaryotic system that can accomplish eukaryotic DNA replication initiation and licensing. The existence of such a system has in fact already been suspected given that: 1) Orc1- or Orc2-depleted human cells and mouse-Orc1 and fruit-fly ORC mutants are viable and capable of undergoing replication and endoreplication^80–83^, and 2) origin-independent replication at the chromosome level has been reported^79, 84, 85^. We propose that *Carpediemonas* species utilize an alternative DNA replication system based on a Dmc1-dependent HR mechanism that is origin-independent and mediated by RNA:DNA hybrids. Here we summarize evidence that such a mechanism is possible based on what is known in model systems and present a hypothetical model as to how it might occur in *Carpediemonas*.

During replication and transcription, the HR complexes, RNAse H1 and RNA-interacting proteins are recruited onto the DNA to assist in its repair^36, 37, 86^. Remarkably, experiments show that HR is able to carry out full genome replication in archaea, bacteria, viruses, and linear mtDNA^85, 87–89^, with replication fork progression rates that are comparable to those of regular replication^90^. A variety of *cis* and *trans* homologous sequences (*e.g.*, chromatids, transcript-RNA or -cDNA) can be used as templates^27, 36, 40^, and their length as well as the presence of one or two homologous ends likely influence a recombination execution checkpoint that decides which HR sub-pathway is utilized^91^. For example, in the absence of a second homologous end, HR by Rad51-dependent break-induced replication (BIR) can either use a newly synthesized DNA strand or independently invade donor sequences, such that the initial strand invasion intermediate creates a migrating D-loop and DNA is synthesized conservatively^27, 91, 92^. Studies have found that BIR does not require the assembly of an ORC complex and Cdc6 but the recruitment of the Cdc7, loading of MCM helicase, firing factors and replicative polymerases are needed for assembling the pre-RC complex^27, 91^. The requirement of MCM for BIR was questioned, as PIF1 helicase was found to be essential for long-range BIR^93^. However, recent evidence shows that MCM is typically recruited for unwinding DNA strands during HR^94, 95^ and is likely needed together with PIF1 to enhance processivity. All these proteins are also suspected to operate during origin-independent transcription-initiated replication (TIR), a still-enigmatic mechanism that is triggered by R-loops resulting from RNA:DNA hybrids during transcription^10, 11, 96^.

Considering the complement of proteins in *Carpediemonas* species discussed above, and that RNA:DNA hybrids are capable of promoting origin-independent replication in model systems^11, 39, 97^, we suggest that a Dmc1-dependent HR replication mechanism is enabled by excess of RNA:DNA hybrids in these organisms. In such a system, DSBs generated in stressed transcription-dependent R-loops could be repaired by HR with either transcript-RNA- or transcript-cDNA-templates and the *de novo* assembly of the replisome as in BIR (**Fig. 4**). The establishment of a replication fork could be favored by the presence of *Carpediemonas*-specific PIF1-like homologs, as these raise the possibility of the assembly of a multimeric PIF1 helicase with increased capability to bind multiple sites on the DNA, thereby facilitating DNA replication processivity and regulation^45^. Note that the foregoing mechanisms will work even if *Carpediemonas* species are haploid as seems likely based on the SNP data. The loss of Rad51A and the duplication of Dmc1 recombinases suggests that a Dmc1-dependent HR mechanism was likely enabled in the last common ancestor of Fornicata and this mechanism may have become the predominant replication pathway in the *Carpediemonas* lineage after its divergence from the other fornicates, ultimately leading to the loss of ORC and Cdc6 proteins.

**Figure 4.**
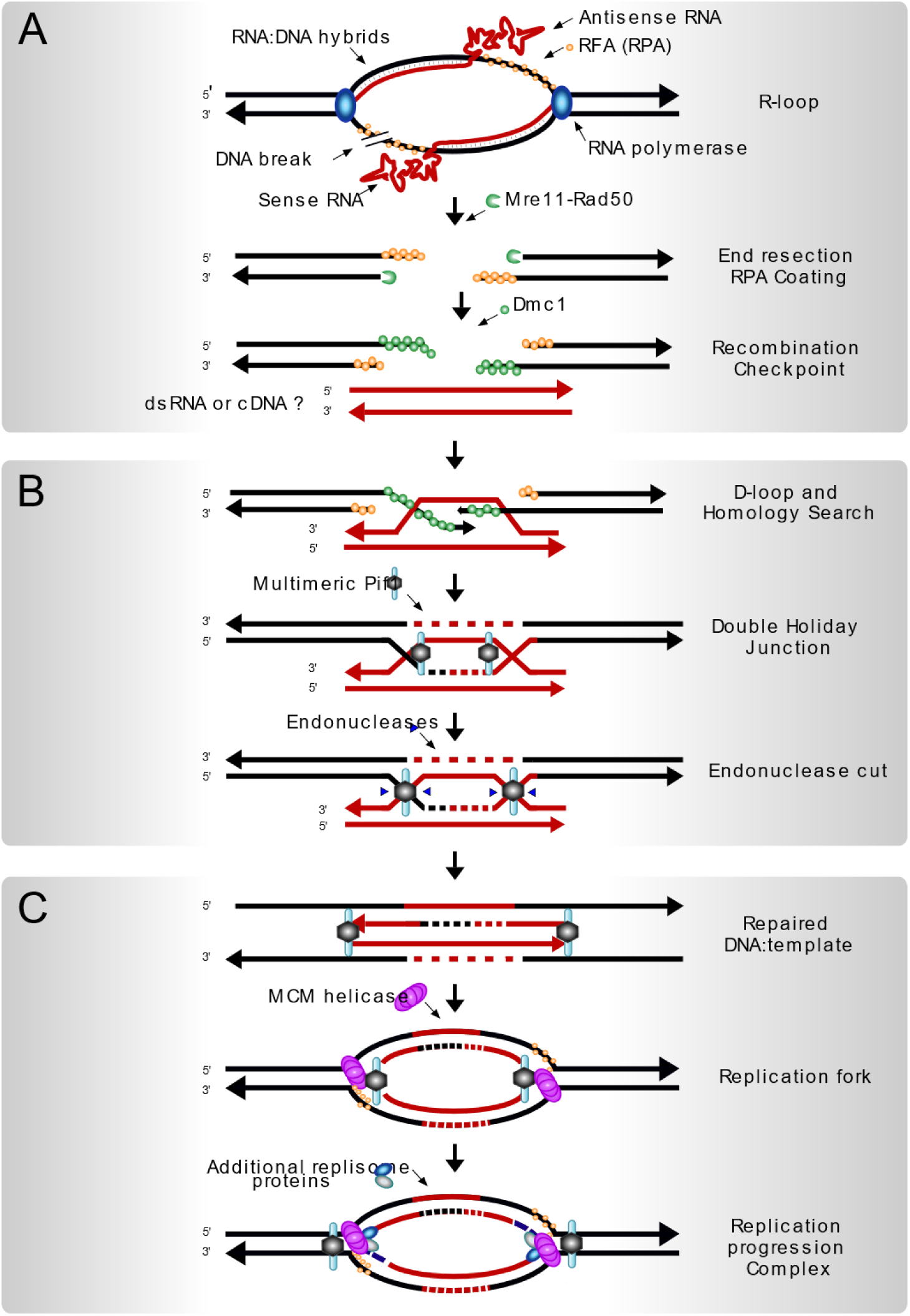
Hypothesis for Dmc1-dependent DNA replication in *Carpediemonas.* **A**) R-loop stimulated sense and antisense transcription^118^ in a highly transcribed locus results in a DNA break, triggering DSB checkpoint control systems to assemble HR complexes and the replication proteins near the lesion^11, 37, 119–121^. Once the damage is processed into a DSB, end resection by Mre11/Rad50 creates a 3’ overhang and the strands are coated with Replication protein A (RPA), while resected ends are coated with the recombinase Dmc1. **B)** A recombination checkpoint decides the HR sub-pathway to be used^91^, then strand invasion of a broken end is initiated into a transcript-RNA or -cDNA template^39, 97, 122^ followed by the initiation and progression of DNA synthesis with the aid of Pif1 helicase*. This leads to the establishment of a double Holliday Junction (HJ) which can be resolved by endonucleases (*e.g.*, Mus81, Flap, Mlh1/Mlh3). The lack of Chk1 may result in mis-segregation caused by aberrant processing of DNA replication intermediates by Mus81^48^. Given the shortness of the RNA or cDNA template, most possible HJ resolutions, except for the one depicted in the figure, would lead to the loss of chromosome fragments. The HJ resolution shown would allow steps shown in panel C. **C)** A multimeric *Carpediemonas* Pif1-like helicase is bound to the repaired DNA as well as to the template. Here, the shortness of the template could resemble a replication intermediate that could prompt the recruitment of MCM, following the addition of the replisome proteins and establishing a fully functional replication fork (Dark blue fragments on 3’ ends of the bottom figure represent Okazaki fragments). *Notes: Polymerases α and δ are able to incorporate the correct nucleotides using RNA template^40^; RNAse H2 would excise ribonucleosides and replace the correct nucleotide.

### The impact of cell cycle dysregulation on genome evolution

DNA replication licensing and firing are temporally separated (*i.e.*, they occur at G1 and S phases respectively) and are the principal ways to counteract damaging over-replication^6^. As S-phase is particularly vulnerable to DNA errors and lesions, its checkpoints are likely more important for preventing genome instability than those of G1, G2 or SAC^98^. Dysregulation is anticipated if no ORC/Cdc6 are present as licensing would not take place and replication would be blocked^28^. Yet this clearly does not happen in *Carpediemonas*. This implies that during late G1 phase, activation by loading the MCM helicase has to occur by an alternative mechanism that is still unknown but might already be in place in eukaryotes. Such a mechanism has long been suspected as it could explain the over-abundance and distribution patterns of MCM on the DNA (*i.e.*, the MCM paradox; reviewed in^99^).

In terms of the regulation of M-phase progression, the extremely divergent nature of the kinetochore in *C. membranifera* could suggests that it uses different mechanisms to execute mitosis and meiosis. It is known that in *Carpediemonas*-related fornicates such as retortamonads and in diplomonads, chromosome segregation proceeds inside a persisting nuclear envelope, with the aid of intranuclear microtubules, but with the mitotic spindle nucleated outside the nucleus (*i.e.*, semi-open mitosis)^78^. Although mitosis in *Carpediemonas* has not been directly observed, these organisms may also possess a semi-open mitotic system such as the ones found in other fornicates. Yet how the *Carpediemonas* kinetochore functions in the complete absence of the microtubule-binding Ndc80 complex remains a mystery; it is possible that, like in kinetoplastids^48^, other molecular complexes have evolved in this lineage that fulfill the roles of Ndc80 and other kinetochore complexes.

Interestingly, a potential repurposing of SAC proteins seems to have occurred in the diplomonad *G. intestinalis*, as it does not arrest under treatment with microtubule-destabilizing drugs and Mad2 localizes to a region of the intracytoplasmic axonemes of the caudal flagella^55^. Other diplomonads have a similar SAC protein complement that may have a similar non-canonical function. In contrast to diplomonads, our investigations (**Fig. 3**) suggest that *Carpediemonas* species could elicit a functional SAC response, although microtubule-disrupting experiments during mitosis will be needed to prove its existence.

In addition to the aforementioned apparent dysregulation of checkpoint controls in *Carpediemonas* species, alternative mechanisms for chromosome condensation, spindle attachment, sister chromatid cohesion, cytokinesis, heterochromatin formation, and silencing and transcriptional regulation can also be expected in this organism due to the absence of ORC and Cdc6 (reviewed in refs^30, 100, 101^). All of the absences of canonical eukaryotic systems we have described for *Carpediemonas* suggest that a radically different cell cycle has evolved in this free-living protistan lineage. This underscores the fact that our concepts of universality and essentiality rely on studies of a very small subset of organisms. The development of *Carpediemonas* as a model system thus has great potential to enhance our understanding of fundamental DNA replication, repair and cell cycle processes. It could even reveal widely conserved alternative, but as-yet unknown, mechanisms underpinning the evolutionary plasticity of these systems across the eukaryote tree of life.

## Methods

### Sequencing, assembly, and protein prediction for *C. membranifera*

DNA and RNA were isolated from log-phase cultures of *C. membranifera* BICM strain (see details in **Supplementary Information**). Sequencing employed Illumina short paired-end and long read (Oxford Nanopore MinION) technologies. For Illumina, extracted, purified DNA and RNA (*i.e.*, cDNA) were sequenced on the Hiseq 2000 (150 x 2 paired-end) at the Genome Québec facility.

Illumina reads were quality trimmed (Q=30) and filtered for length (>40 bp) with Trimmomatic^102^. For MinION, the library was prepared using the 1D native barcoding genomic DNA (SQK-LSK108 with EXP-NBD103) protocol (NBE_9006_v103_revP_21Dec2016). The final library (1070 ng) was loaded on a R9.4 flow cell and sequenced for 48 h on the MinION Mk1B nanopore sequencer. The long reads were base-called and trimmed with Albacore v2.3.3 (www.nanoporetech.com) and Porechop v0.2.3 (www.github.com/rrwick/Porechop), respectively. ABruijn v1.0 (www.github.com/fenderglass/Flye/releases/tag/1.0) with default parameters and max genome size of 30Mb produced an assembly that was polished with Nanopolish v0.10.1^103^. The latter was iteratively error-corrected with the genomic paired-end Illumina reads using Unicycler^104^. The identification and removal of prokaryotic contigs was assisted by BLASTn searches against the nt database. Read-depth coverage at each position of the genomic scaffolds were obtained with samtools^105^ and mosdepth v0.2.5^106^.

RNA-Seq reads were used for genome-independent assessments of the presence of the proteins of interest and to generate intron junction hints for gene prediction. For the independent assessments we obtained both a *de novo* and a genome-guided transcriptome assembly with Trinity v2.5.0^107^. Open reading frames were translated with TransDecoder v5.5.0 (www.github.com/TransDecoder) and were included in all of our analyses. Gene predictions were carried out as follows: repeat libraries were obtained and masked with RepeatModeler 1.0 and RepeatMasker (http://www.repeatmasker.org).

Then, RNA-Seq reads were mapped onto the assembly using Hisat2^108^, generating a bam file for GenMarkET^109^. This resulted in a list of intron hints used to train Augustus v3.2.3^110^. The genome-guided assembled transcriptome, genomic scaffolds and the newly predicted proteome were fed into the PASA pipeline^111^ to yield a more accurate set of predicted proteins. Finally, the predicted proteome was manually curated for the proteins of interest.

### Genome size, completeness, and ploidy assessments

We estimated the completeness of the draft genome by 1) using the k-mer based and reference free method Merqury^23^, 2) calculating the percentage of transcripts that aligned to the genome, and 3) employing the BUSCO^112^ framework. For method 1, all paired-end reads were used to estimate the best k-mer and create ‘meryl’ databases necessary to apply Merqury^23^. For method 2, transcripts were mapped onto the genome using BLASTn and exonerate^113^. For method 3, the completeness of the draft genome was evaluated in a comparative setting by including the metamonads and using the universal single copy orthologs (BUSCO) from the Eukaryota (odb9) and protist databases (https://busco.ezlab.org/), which contain 303 and 215 proteins, respectively. Each search was run separately on the assembly and the predicted proteome for all these taxa. Unfortunately, both BUSCO database searches yielded false negatives in that several conserved proteins publicly reported for *T. vaginalis*, *G. intestinalis* and *Spironucleus salmonicida* were not detected due to the extreme divergence of metamonad homologs. Therefore, genome completeness was re-assessed with a phylogeny-guided search (**Supplementary Information**).

The ploidy of *C. membranifera* was inferred by *i*) counting k-mers with Merqury^23^, and *ii*) mapping 613,266,290 Illumina short reads to the assembly with Bowtie 2.3.1^114^ and then using ploidyNGS^115^ to calculate the distribution of allele frequencies across the genome. A site was deemed to be heterozygous if at least two different bases were present and there were at least two reads with the different bases. Positions with less than 10× coverage were ignored.

### Functional annotation of the predicted proteins

Our analyses included the genomes and predicted proteomes of *C. membranifera* (reported here) as well as publicly available data for nine additional metamonads and eight other eukaryotes representing diverse groups across the eukaryotic tree of life (**Fig. 1**, **Supplementary Information**). Orthologs from each of these 18 predicted proteomes were retrieved for the assessment of core cellular pathways, such as DNA replication and repair, mitosis and meiosis and cell cycle checkpoints. For *C. membranifera,* we included the predicted proteomes derived from the assembly plus the 6-frame translated transcriptomes. Positive hits were manually curated in the *C. membranifera* draft genome. A total of 367 protein queries were selected based on an extensive literature review and prioritizing queries from taxa in which they had been experimentally characterized. The identification of orthologs was as described for the BUSCO proteins but using these 367 queries for the initial BLASTp (**Supplementary Information**), except for kinetochore (KT), Spindle assembly check point (SAC) and anaphase-promoting complex-related genes (APC/C). For these, previously published refined HMMs with cut-offs specific to each orthologous group (see^58^) were used to query the proteomes with HMMER v3.1b2^32^. A multiple sequence alignment that included the newly-found hits was subsequently constructed with MAFFT v7.310^116^ and was used in HMM searches for more divergent homologs. This process was iterated until no new significant hits could be found. As we were unable to retrieve orthologs of a number of essential proteins in the *C. membranifera* and *C. frisia* genomes, we embarked on additional more sensitive strategies to detect them using multiple different HMMs based on aligned homologs from archaea, metamonads, and broad samplings of taxa. Individual PFAM domains were searched for in the genomes, proteome and transcriptomes with e-value thresholds of 10^-3^ (**Supplementary Information**). To rule out that failure to detect these proteins was due to insufficient sensitivity of our methods when applied them to highly divergent taxa, we queried 22 extra eukaryotic genomes with demonstrated high rates of sequence evolution, genome streamlining or unusual genomic features (**Supplementary Table 1**, **Supplementary Information**). Possible non-predicted or mis-predicted genes were investigated using tBLASTn searches of the genomic scaffolds and unassembled reads and 6-frame translation searches with HMMER. Also, as DNA replication and repair genes could have been acquired by lateral gene transfer into *Carpediemonas* species from prokaryotes or viruses, proteins from the DNA replication and repair categories whose best matches were to prokaryotic and viral homologs were subjected to phylogenetic analysis using the methods described for the phylogeny-guided BUSCO analysis and using substitution models specified in the legend of each tree (**Supplementary Information**).

### Data availability

Genome assembly will be available at NCBI under BioProject PRJNA719540, biosample number SAMN18612951, accession numbers <XXXX>.

## Acknowledgments

The majority of this work was supported by a Foundation grant FRN-142349, awarded to A.J.R. by the Canadian Institutes of Health Research. Archibald Lab contributions to this study were supported by a Discovery Grant from the Natural Sciences and Engineering Research Council of Canada (RGPIN 05871-2014). E.C.T. is supported by a Herchel Smith Postdoctoral Fellowship at the University of Cambridge.

## Author contributions

D.E.S-L and A.J.R. conceived the study. J.J-H and M.K. grew cultures, extracted nucleic acids, and carried out *in house* sequencing. D.E.S-L., B.A.C., E.C.T., Z.Y, J.S.S-L., L.G-L., G.J.P.L.K, J.M.A., A.G.B.S. and A.J.R. analyzed and manually curated the genomic data. E.C.T. and D.E.S-L made the figures. D.E.S-L and A.J.R. led the writing of the manuscript with input from all authors. All documents were edited and approved by all authors.

## Additional information

Supplementary Information (also containing legends for Supplementary Table 1 and Supplementary Figures 1 – 10)

## SUPPLEMENTARY INFORMATION

### A. Supplementary methods

#### A1. Culturing and DNA isolation

Sequencing of *C. membranifera* BICM strain was done with Illumina short paired-end and long MinION read technologies. The Illumina sequencing employed DNA from a monoxenic culture grown in 50 ml Falcon tubes in F/2 media enriched with the bacterium *Shewanella frigidimarina* as food. DNA was isolated from a total of two litres of culture using a salt extraction protocol followed by CsCl gradient centrifugation. RNA was also extracted from these cultures using TRIzol (Invitrogen, USA), following the manufacturer’s instructions. For MinION sequencing, *C. membranifera* was grown in sterile filtered 50% natural sea water media with 3% LB with either *Shewanella sp* or *Vibrio sp.* isolate JH43 as food. Cell cultures were harvested at peak density by centrifugation at 500×*g*, 8 min, 20 °C. The cells were resuspended in sterile-filtered spent growth media (SFSGM) and centrifuged again at 500×*g*, 8 min, 20 °C. The cell pellets were resuspended in 1.5 mL SFSGM, layered on top of 9 mL Histopaque®-1077 (Sigma-Aldrich) and centrifuged at 2000×*g*, 20 min, 20 °C. The protists were recovered from the media:Histopaque interface by pipetting, diluted in 10 volumes of SFSGM and centrifuged 500×*g*, 8 min, 20 °C. High molecular weight DNA was extracted using MagAttract HMW DNA Kit (Qiagen, Cat No. 67563), purified with GenomicTip 20/G (Qiagen, Cat No. 10223) and resuspended in 5 mM Tris-HCl (pH 8.5).

#### A2. Genome size and completeness using BUSCO and a phylogeny-guided approach

The BUSCO approach^1^ was prone to false negative predictions with our dataset because of the extreme divergence of metamonad homologs. Therefore, the completeness of the BUSCO set was re-assessed with a phylogeny-guided search. For this, we eliminated 31 proteins associated with mitochondria or mitochondrion-related organelles (MROs) as Metamonada have reduced or no MROs^2^, and employed taxa-enriched Hidden Markov Model (HMM) searches to account for divergence between the remaining 272 proteins and the studied taxa. In brief: BLASTp was carried out using the 272 BUSCO proteins as queries for finding their orthologues in a local version of the PANTHER 14.0 database^3^ to enable the identification of the most likely Panther subfamily HMM and its annotation. Then, each corresponding subfamily HMM was searched for in the predicted proteomes with an e-value cut-off of 1×10^-1^ with HMMER v3.1b2^4^. In cases where these searches did not produce any result, a broader search was run using the HMM of the Panther family with 1×10^-3^ as e-value cut-off. Five best hits for each search were retrieved from each proteome, aligned to the corresponding Panther subfamily or family sequences with MAFFT v7.310^5^ and phylogenetic reconstructions were carried out using IQ-TREE v1.6.5^6^ under the LG+C60+F+Γ model with ultrafast bootstrapping (1000 replicates). Protein domain architectures were visualized by mapping the respective Pfam accessions onto trees using ETE tools v3.1.1^7^.

#### A3. Taxa selected for comparative genomic analysis

Our analyses included the publicly available genomes and predicted proteomes of *Trichomonas vaginalis* G3 (Parabasalia, www.trichdb.org), *Monocercomonoides exilis* (Preaxostyla, www.protistologie.cz/hampllab), the free-living fornicates *Carpediemonas frisia*^8^ (*i.e.*, metagenomic bin and predicted proteome), *Carpediemonas membranifera* (reported here) and *Kipferlia bialata*^9^, plus the parasitic diplomonad fornicates: *Giardia intestinalis* Assemblages A and B, *Giardia muris*, *Spironucleus salmonicida*-ATCC50377 (www.giardiadb.org) and *Trepomonas* PC1^10^ −the latter was only available as a transcriptome. We also included a set of genomes that are broadly representative of eukaryote diversity, such as *Homo sapiens* GRCh38, *Saccharomyces cerevisiae* S288C, *Arabidopsis thaliana* TAIR10, *Dictyostelium discoideum* AX4, *Trypanosoma brucei* TREU927-rel28 (www.uniprot.org), *Naegleria gruberi* NEG-M (www.ncbi.nlm.nih.gov), *Guillardia theta* and *Bigelowiella natans* (www.genome.jgi.doe.gov/portal/).

Additional analyzed genomes were those of the microsporidia *Encephalitozoon intestinalis* ATCC 50506 (ASM14646v1), *E. cuniculi* GB-M1 (ASM9122v2) and *Trachipleistophora hominis* (ASM31613v1), the yeasts *Hanseniaspora guilliermondii* (ASM491977v1), *Hanseniaspora opuntiae* (ASM174979v1), *Hanseniaspora osmophila* (ASM174704v1), *Hanseniaspora uvarum* (ASM174705v1) and *Hanseniaspora valbyensis* NRRL Y-1626 ( GCA_001664025.1), *Tritrichomonas foetus* (ASM183968v1), the nucleomorphs of *Hemiselmis andersenii* (ASM1864v1), *Cryptomonas paramecium* (ASM19445v1), *Chroomonas mesostigsmatica* (ASM28609v1), *Guillardia theta* (ASM297v1), *Lotharella vacuolata* (AB996599–AB996601), *Amorphochlora amoebiformis* (AB996602–AB996604) and *Bigellowiela natans* (ASM245v1), the corals *Galaxea fascicularis*, *Fungia sp*., *Goniastrea aspera*, *Acropora tenuis* and the coral endosymbionts *Symbiodinium kawagutii* and *Symbiodinium goreaui*^11, 12^.

#### A4. Additional strategies used to search for ORC, Cdc6 ad Ndc80 proteins

Strategies included enriched HMMs as mentioned in the main text and HMMs for individual Pfam domains with e-value thresholds of 1×10^-3^. 1) Metamonad-specific HMMs were built as described for kinetochore proteins − containing the newly found hits plus orthologs from additional publicly available metamonad proteomes or transcriptomes^2, 13^, 2) we applied the eggNOG 4.5 profiles COG1474, COG5575, KOG2538, KOG2228, KOG2543, KOG4557, KOG4762, KOG0995, KOG4438, KOG4657 and 2S26V which encompass 2774, 495, 452, 466, 464, 225, 383, 504, 515, 403 and 84 taxa, respectively, and 3) the Pfam v33.1 HMMs: PF09079 (Cdc6_C), PF17872 (AAA_lid_10), PF00004 (AAA+), PF13401 (AAA_22), PF13191 (AAA_16), PF01426 (BAH), PF04084 (Orc2), PF07034 (Orc3), PF18137 (ORC_WH_C), PF14629 (Orc4_C), PF14630 (Orc5_C), PF05460 (Orc6), PF03801 (Ndc80_HEC), PF03800 (Nuf2), PF08234 (Spindle_Spc25) and PF08286 (Spc24). For Ncd80, Nuf2, Spc24 and Spc25 we also applied the HMMs models published in^14^.

### B. Supplementary results

#### B1. BUSCO completeness

A subset of 272 BUSCO proteins from the odb9 database was used for a phylogeny-guided search for divergent orthologs. This revealed that: *i*) 27 out of 272 BUSCO (9.9%) proteins are absent in all metamonads, *ii*) only 101 (∼41%) of the remaining 245 proteins were shared by all metamonad proteomes, and *iii*) up to 38% are absent in all Fornicata. Metamonad genomes only contained 60% to 91% of the BUSCO proteins (**Table 1**, **Supplementary Table 1**, note that the BUSCO presence-absence patterns of the transcriptomic data from *Trepomonas* sp. PC1 are consistent with those of the remaining diplomonads). These analyses demonstrate that the Metamonada have secondarily lost a relatively large number of highly conserved eukaryotic proteins and, therefore, BUSCO analysis cannot be used on its own to evaluate metamonad genome completeness.

#### B2. Additional search strategies to find missing proteins

Metamonad-specific HMM retrieved two candidates for Orc1/Cdc6 proteins from *C. frisia* (*i.e.*, Cfrisia_2222, Cfrisia_2845) and one from *C. membranifera* (*i.e.*, c4603.t1), and one Orc4 candidate from each *Carpediemonas* species (*i.e.*, Cfrisia_2559, ds58_16707). Further inspection of these hits showed that only the AAA+ region shared similarity among all of these proteins, which is expected as ORC and Cdc6 proteins belong to the ATPase superfamily. However, based on full protein identity, full profile composition and domain architecture, the proteins retrieved with the Orc1/Cdc6 HMM were confidently annotated as Katanin P60 ATPase-containing subunit A1 (Cfrisia_2222), Replication factor C subunits 1 (c4603.t1) and 5 (Cfrisia_2845), and proteins retrieved with Orc4 HMM were members of the Dynein heavy chain (Cfrisia_2559) and AAA-family ATPase families (ds58_16707). The latter is a 744 aa protein that has a C-terminal region with no sequence similarity or amino acid profile frequencies that resembles a Orc4_C Pfam domain from other metamonads or model eukaryotes. All the additional search strategies yielded false positives in *Carpediemonas* species, as these retrieved AAA-family members lacking sequence similarity to orc proteins, showed completely different protein domain architecture than the expected one and were associated with different functional annotation (data not shown). When reconstructing the domain architecture of ORC and Cdc6 proteins in metamonads, we noted that Fornicata Orc1/Cdc6-like proteins are remarkably smaller (*i.e.*, 1.5 to 3 times smaller) than Orc1 and Cdc6 from the model organisms and other protists used later in phylogenetic reconstruction (**Supplementary Figure 1A and B, Supplementary Table 1**). In most cases, the small proteins lack protein domains rendering a different domain architecture with respect to their homologs in *S. cerevisiae*, *H. sapiens, A. thaliana* and *T. vaginalis* (**Supplementary Figure 1A**, **Supplementary Table 1**). For example, Orc1 and Cdc6 paralogs in Fornicata lack BAH, and AAA_lid10 and Cdc6_C domains. Protein alignments show that the conserved areas of these proteins correspond to AAA+ domain that have relatively conserved Walker domains A and B (except MONOS_13325 from *M. exilis*), with a few proteins lacking the arginine finger motif (R-finger) within the Walker B motif (**Supplementary Figure 1B**). The latter may negatively affect ATPase activity of the R-finger-less proteins. In an attempt to establish orthology, metamonad Orc1/Cdc6 candidates were used for phylogenetic reconstruction together with publicly available proteins that have reliable annotations for Orc1 and Cdc6, expected domain architecture and/or with experimental evidence of their functional activity in the replisome. Phylogenetic analysis shows that metamonad proteins form separate clades from the *bona fide* Orc1 and Cdc6 sequences (**Supplementary Figure 1C**). One of these separate clades encompasses Orc1-b from *T. brucei* that has been shown to participate during DNA replication despite lacking the typical domain architecture^15^.

#### B3. DNA replication streamlining in nucleomorphs

The loss of ORC/Cdc6 accompanied by the partial retention of MCM, PCNA, Cdc45, RCF, GINS and the homologous recombination (HR) recombinase Rad51 was observed in cryptophyte and chlorarachniophyte nucleomorphs (**Supplementary table 1**). ORC and Cdc6 were found as single copies (except Orc2) in the nuclear DNA of these two groups; their predicted proteins lack obvious signal and targeting peptides which would likely prevent them from participating in a nucleus-coordinated nucleomorph replication. Hence, nucleomorph DNA replication likely occurs by HR without the assistance of ORC/Cdc6 origin-binding, but this replication might nonetheless be regulated at the transcriptional level by the nucleus as shown by^16^. Many of the remaining nuclear-encoded proteins involved in replication are present in more than one copy in those taxa, with several of them containing signal and transit peptides (*e.g.*, H2A, POLD, RCF1 and RFA1)^16, 17^.

#### B4. Acquisition of Endonuclease IV, RarA and RNAse H1 by lateral gene transfer

The Endonuclease IV (Apn1 in yeast) and exonuclease III (Exo III) function in the removal of abasic sites in DNA via the BER pathway. Our analyses show that *C. frisia* and *C. membranifera* have Exo III and have a prokaryotic version of Endo IV (**Supplementary Fig 8**). Interestingly, none of the parabasalids and *Giardia* spp. have an Endo IV homolog, either eukaryotic or prokaryotic. *S. salmonicida* and *Trepomonas* sp. PC1, by contrast, appear to encode a typical eukaryotic Endo IV.

The RarA (Replication-Associated Recombination protein A, also named MgsA) protein is ubiquitous in bacteria and eukaryotes (*e.g.*, homologs Msg1 in yeast and WRNIP1 in mammals) and acts in the context of collapsed replication forks^18, 19^. *Carpediemonas* possesses a prokaryotic-like version (**Supplementary Fig 9**) that lacks the ubiquitin-binding Zn finger N-terminal domain typical of eukaryotic homologs^18^. No canonical eukaryotic RarAs were detected in the remaining metamonads, but it appears that prokaryotic-like RarA proteins in *Giardia*, *S. salmonicida* and *Trepomonas* sp. PC1 were acquired in an independent event from that of *Carpediemonas*.

Both *Carpediemonas* genomes have a eukaryotic RNAse H2, lack eukaryotic RNAse H1 but encode up to two copies of a prokaryotic-like RNAse H1 (**Supplementary Fig. 10**) which do not have the typical eukaryotic HBD domain^20^. The HBD domain is thought to be responsible for the higher affinity of this protein for DNA/RNA duplexes rather than for dsRNA^21, 22^. All prokaryotic-like RNAse H1s in metamonads are highly divergent (**Supplementary Fig. 10**) and, in the case of *S. salmonicida* RNaseH1 proteins, these formed very long branches in all of our preliminary trees, that had to be removed for the final phylogenetic reconstruction. Remarkably, the phylogenetic reconstruction that includes other metamonad proteins suggests that *Giardia*, *Trepomonas* sp. PC1, *T. foetus* and *T. vaginalis*, also acquired bacterial RNAseH1. *Trepomonas sp*. PC1 and *Giardia* sequences cluster together but the *T. foetus* and *T. vaginalis* enzymes each emerge amidst different bacterial branches, suggesting that they have been acquired independently from the *Carpediemonas* homologs. It should, however, be noted that the support values are overall low, partly due to the fact that these sequences and their relatives are highly divergent from each other, from *Carpediemonas* bacterial-like sequences, and from typical eukaryotic RNaseH1.

### C. Supplementary discussion

#### C1. BUSCO incompleteness

Both eukaryote-wide and protist BUSCO analyses using the BUSCO methods underperformed in our analyses. Despite using a phylogeny-guided search with the Eukaryota database, a more comprehensive database than the protist BUSCO database, a remarkably large number of BUSCO proteins were inconsistently present in Metamonada. This is not surprising, as the clade harbors a very diverse group of taxa with varied lifestyles and many have undergone genome streamlining^9, 10, 23–25^, and the BUSCO databases are expected to be more accurate with greater taxonomic proximity to the studied genome^1, 26, 27^. While it might be tempting to suggest the 101 BUSCO proteins that are shared by all metamonads be used to evaluate genome completion in the clade, the overwhelming evidence of differential genome streamlining strongly indicates that databases should be lineage specific (*e.g.*, *Carpediemonas*, *Giardia*, etc). Hence, our results highlight the need for constructing such databases including proteins that showcase the sequence diversity of the groups and genes that are truly single copy in each of these lineages. Regardless, using only standard BUSCO methods to capture genome completion will still fall short in such assessments as it will fail to evaluate the most difficult-to-assemble regions of the genome^27, 28^. For that reason, combined approaches such as the ones used here provide a more comprehensive global overview of genome completeness.

## E.

**Supplementary figures (Note:** 1166 Any reference in Supplementary Figure legends can be found in Supplementary References)

**Supplementary Fig 1.**
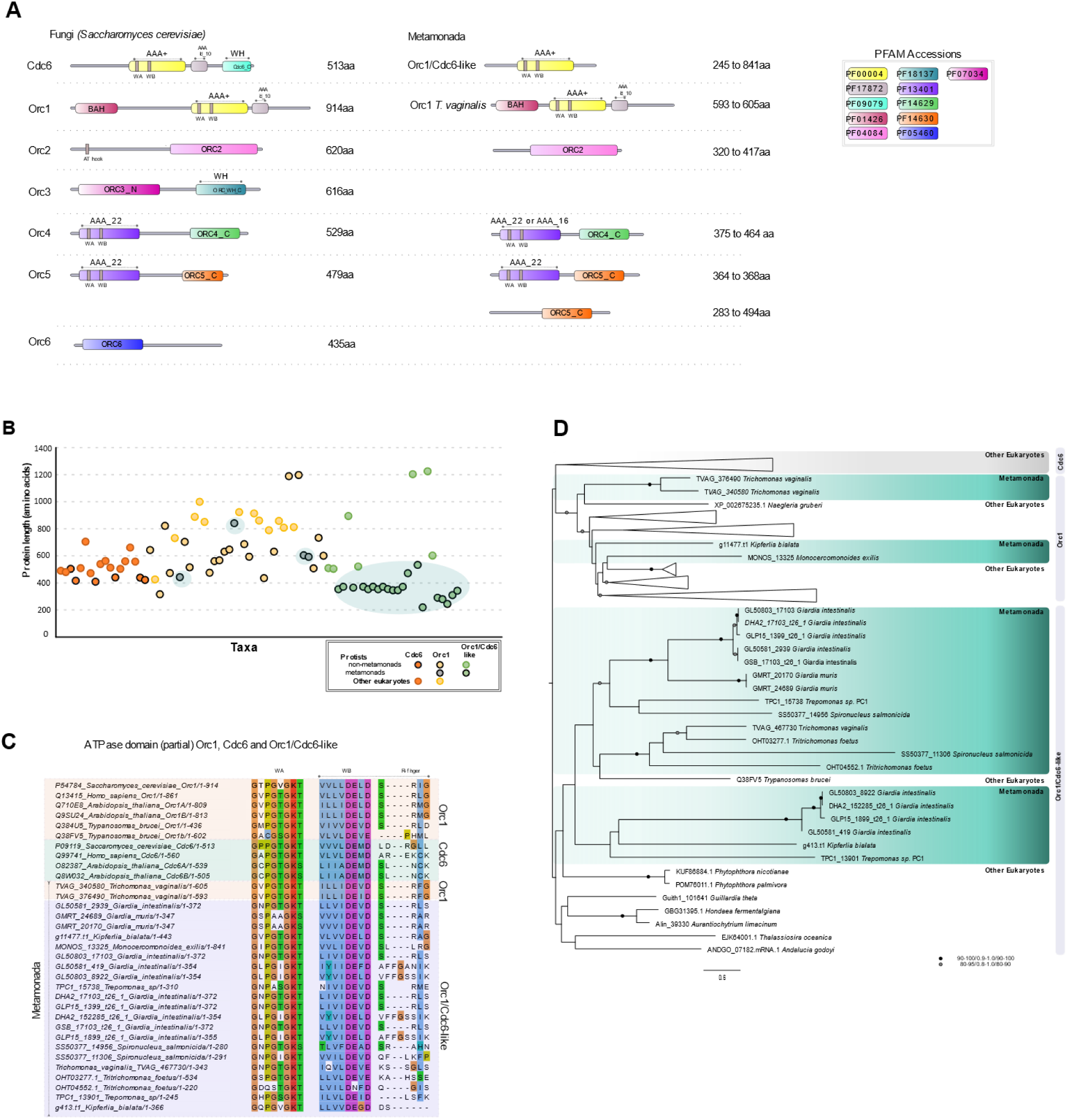
Orc1-6 and Cdc6 proteins. **A)** Left: typical domain architecture observed for Orc1-6 and Cdc6 in *Saccharomyces cerevisiae*, Right: representative domain architecture of metamonad proteins drawn to reflect the most common protein size. If no species name is given, then the depicted domain structure was found in all of the metamonads where present. Numbers on the right of each depiction correspond to the total protein length or its range in the case of metamonads (additional information in **Supplementary** Table 1). B) Comparison of Orc1, Cdc6 and Orc1/Cdc6-like protein lengths across 81 eukaryotes encompassing metamonads and non-metamonads protists (source information in **Supplementary** Table 1). Metamonad proteins are highlighted with green shaded bubbles in the background. **C)** Orc1/Cdc6 partial ATPase domain showing Walker A and Walker B motifs including R-finger. Reference species at the top. Multiple sequence alignment was visualized with Jalview^29^ using the Clustal colouring scheme. **D)** Phylogenetic reconstruction of Orc1, Cdc6 and Orc1/Cdc6-like proteins inferred with IQ-TREE^6^ under the LG+ C10+F+ Γ model using 1000 ultrafast bootstraps (bootstrap value ranges for branches are shown with black and grey dots). The alignment consists of 81 taxa with 367 sites after trimming. Orc1/Cdc6-like proteins do not form a clade with *bona fide* Orc1 and Cdc6 proteins making it impossible to definitively establish whether or not they are orthologs.

**Supplementary Fig 2.**
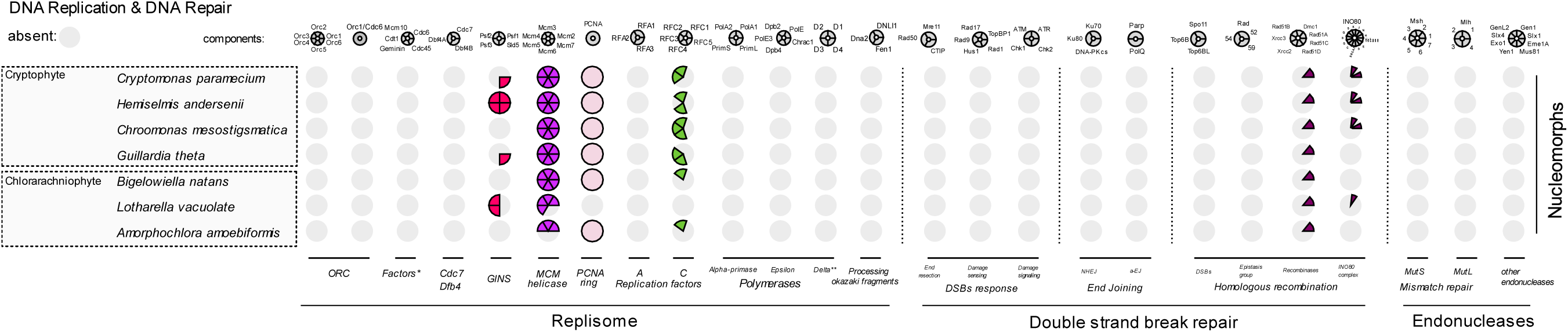
The distribution of core molecular systems of the replisome, double strand break repair and endonucleases in nucleomorph genomes of cryptophyte and chlorarachniophytes.

**Supplementary Fig 3.**
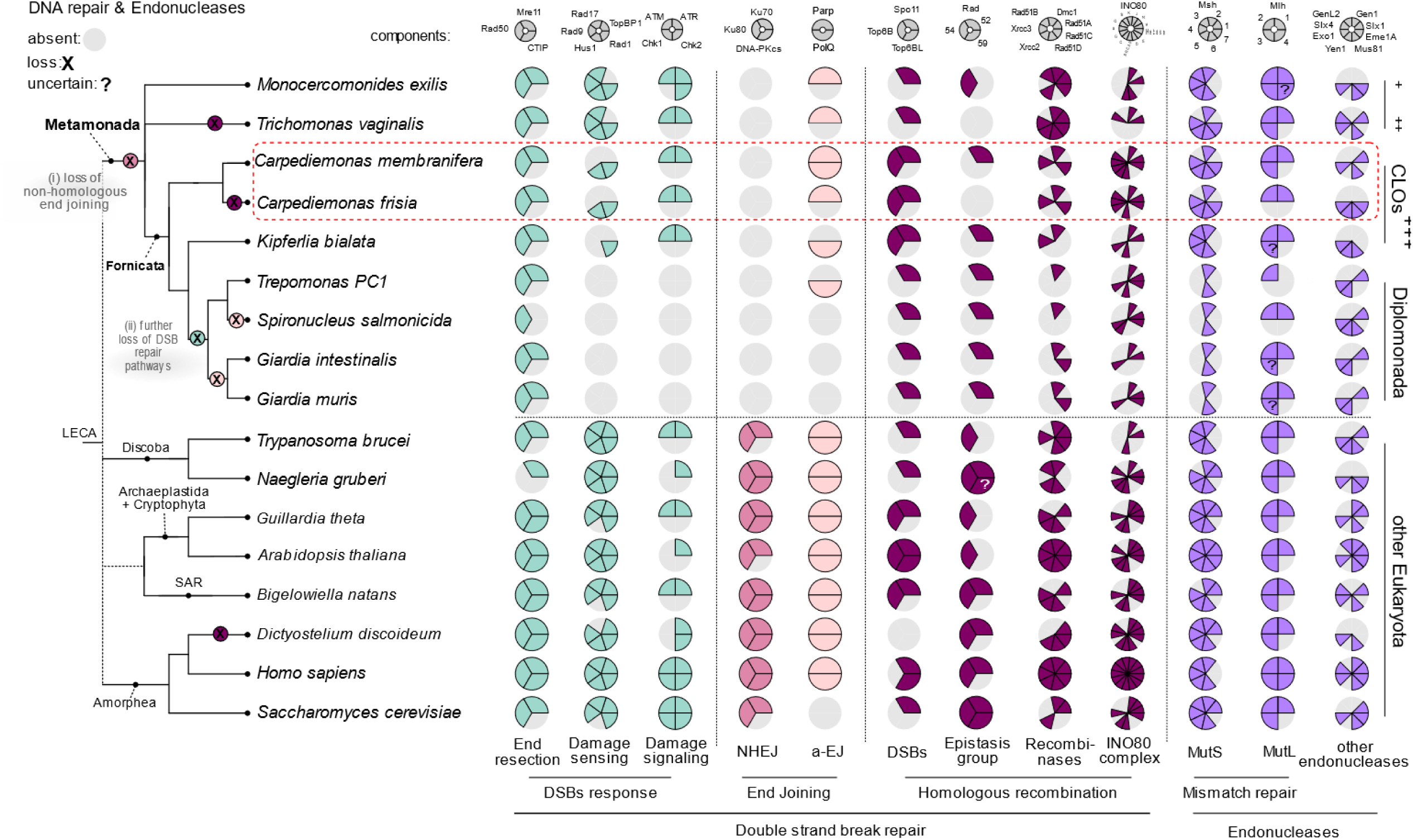
The distribution of core molecular systems of DNA repair across eukaryotic diversity. A schematic global eukaryote phylogeny is shown on the left with classification of the major metamonad lineages indicated. Double strand break repair and endonuclease sets. \*\*\**Carpediemonas*-Like Organisms. ‘?’ is used in cases where correct orthology was difficult to establish, so the protein name appears with the suffix ‘-like’ in tables.

**Supplementary Fig 4.**
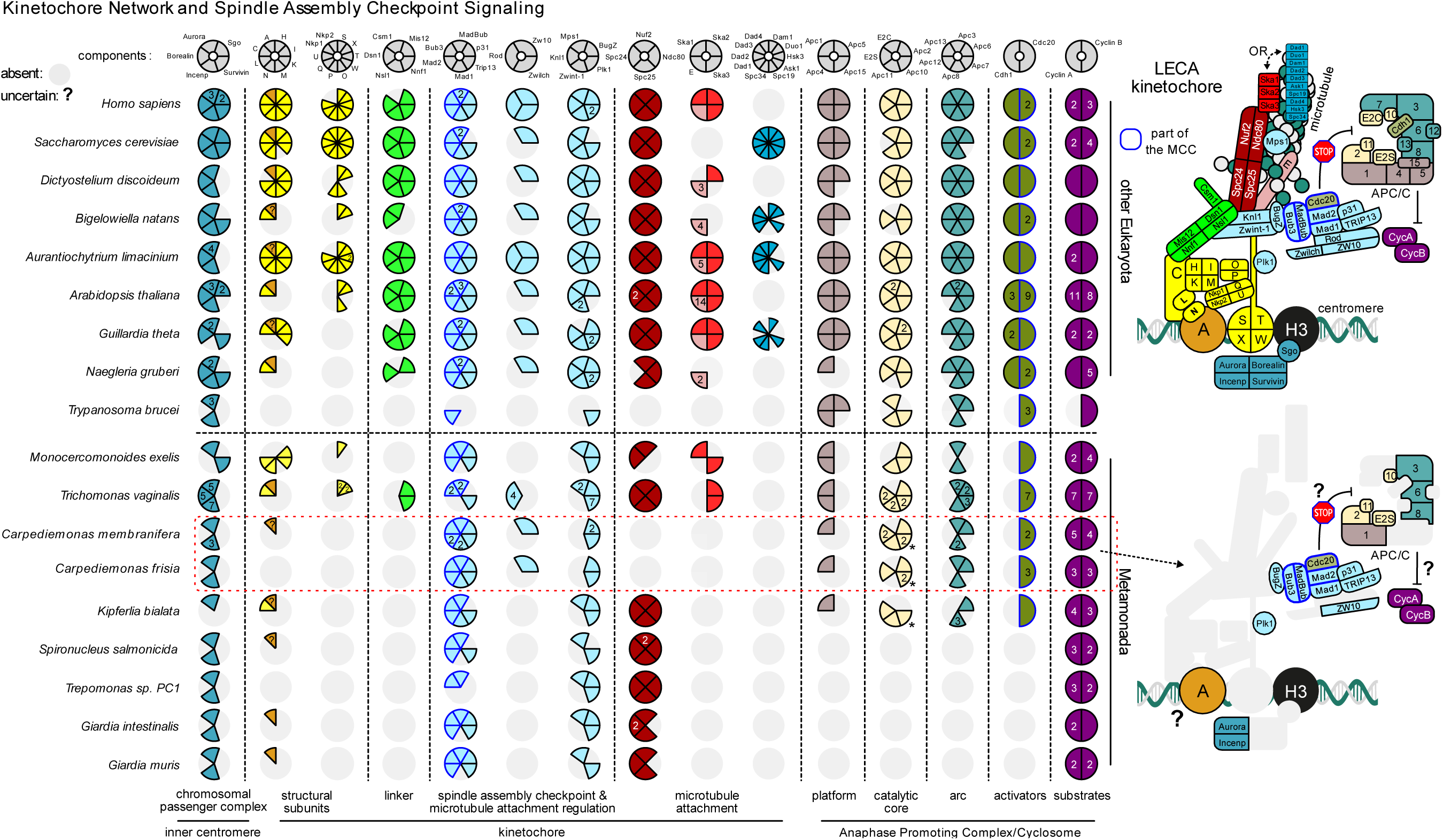
Presence/absence diagram of LECA kinetochore components in eukaryotes, with a greater sampling of metamonads, including *C. membranifera* and *C. frisia*. Left: matrix of presences (coloured) and absences (light grey) of kinetochore, SAC and APC/C proteins that were present in LECA. On top: names of the different subunits; single letters (A-X) indicate Centromere protein A-X (*e.g*., CenpA) and numbers, APC/C subunit 1-15 (*e.g.*, Apc1). E2S and E2C, refer to E2 ubiquitin conjugases S and C, respectively. Colour schemes correspond to the kinetochore overview figure on the right and to that used in Figure 1. Right: cartoon of the components of the kinetochore, SAC signalling, the APC/C and its substrates (Cyclin A/B) in LECA and Carpediemonas species to indicate the loss of components (light grey shading). Blue lines indicate the presence of proteins that are part of the MCC. Asterisk: Apc10 has three paralogs in *C. membranifera* and two in *C. frisia*. One is the canonical Apc10, the two others are fused to a BTB-Kelch protein of which its closest homologs is a likely adapter for the E3 ubiquitin ligase Cullin 3.

**Supplementary Fig 5.**
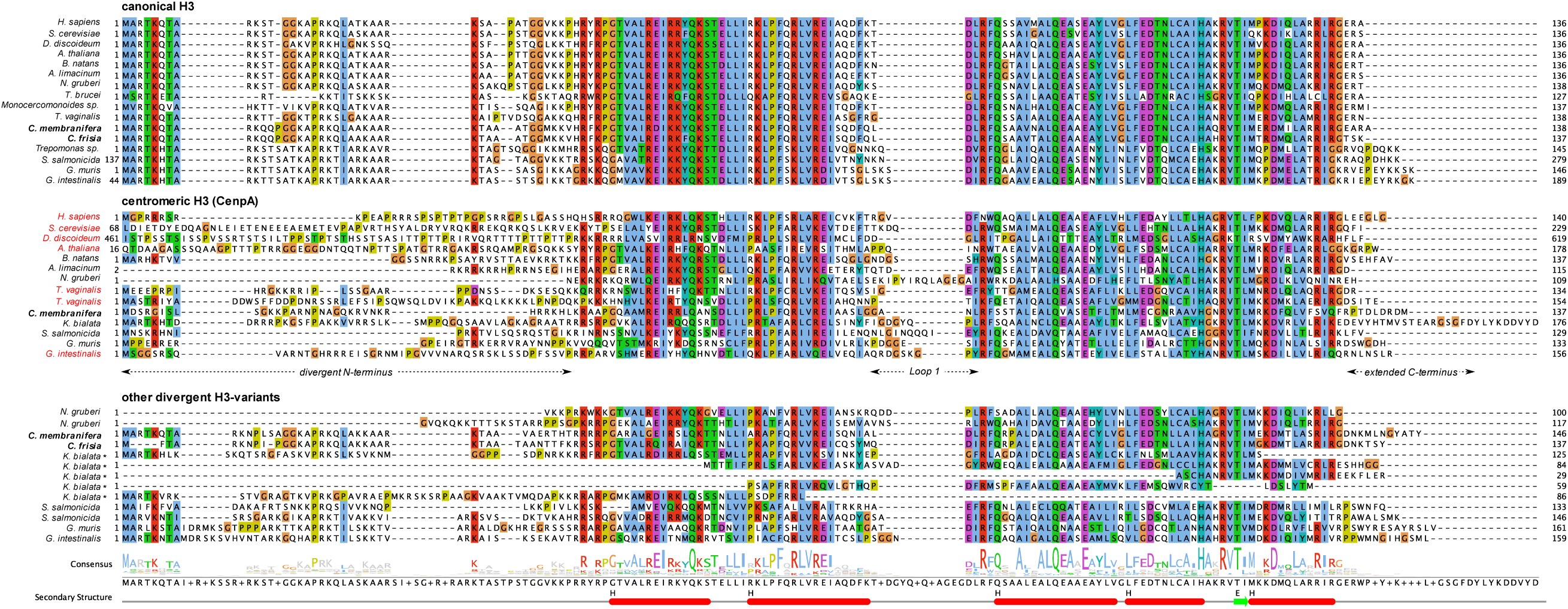
*Carpediemonas* harbours three different types of Histone H3 proteins, a centromere-specific variant (CenpA). Multiple sequence alignment of different Histone H3 variants in eukaryotes and metamonads, including the secondary structure of canonical H3 in humans (pdb: 6ESF_A). CenpA orthologs are characterized by extended amino and carboxy termini and a large L1 loop. Red names in the CenpA panel indicate for which species centromere/kinetochore localization has been confirmed. In addition to CenpA and canonical Histone H3-variants, multiple eukaryotes, including *C. membranifera* and *C. frisia*, harbour other divergent H3 variants. Such divergent variants make the annotation of Histone H3 homologs ambiguous (see Asterisks; incomplete sequences). Multiple sequence alignments were visualized with Jalview^29^, using the Clustal colour scheme. Asterisks indicate two potential CenpA candidates in *T. vaginalis*.

**Supplementary Fig 6.**
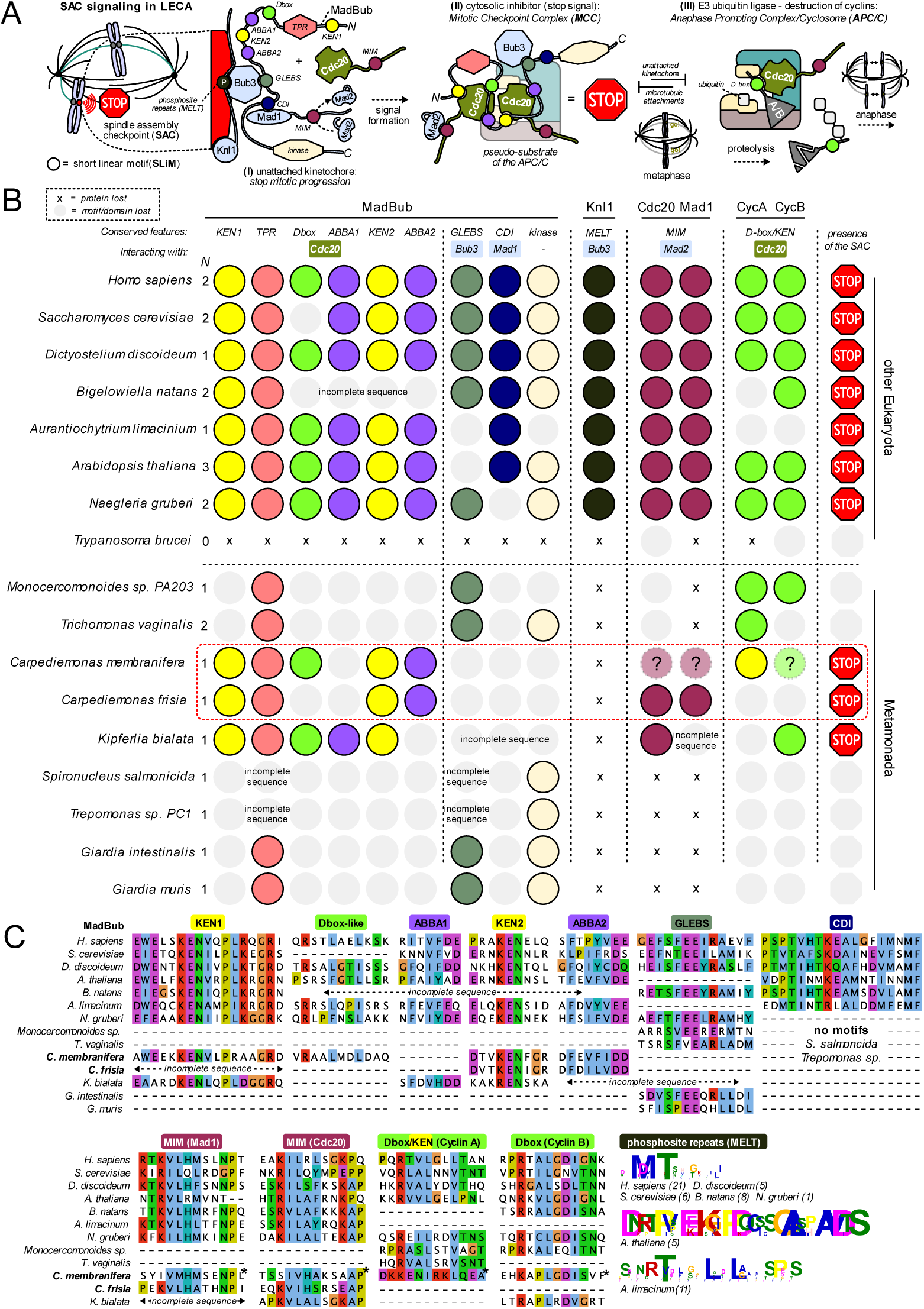
Likely presence of SAC signalling in *Carpediemonas*. A) Short linear motifs form the basis of SAC signalling. During prometaphase, unattached kinetochores catalyse the production of inhibitor of the cell cycle machinery, a phenomenon known as the SAC^30^. **(I)** The main protein scaffold of SAC signalling is the kinase MadBub (paralogs Mad3/Bub1 exist in eukaryotes), which consist of many short linear motifs (SLiMs) that mediate the interaction of SAC components and the APC/C (light blue)^31, 32^. MadBub itself is recruited to the kinetochore through interaction with Bub3 (GLEBS), which on its turn binds repeated phosphomotifs in Knl1^33–35^. The CDI or CMI motif aids to recruit Mad1^36–38^, which has a Mad2-interaction Motif (MIM) that mediated the kinetochore-dependent conversion of open-Mad2 to Mad2 in a closed conformation^39^. **(II)** Mad2, MadBub, Bub3 and 2x Cdc20 (APC/C co-activator) form the mitotic checkpoint complex (MCC) and block the APC/C^32, 40, 41^. MadBub contains 3 different APC/C degrons (D-box, KEN-box and ABBA motif)^31^ that direct its interaction with 2x Cdc20s and effectively make the MCC a pseudo substrate of the APC/C. **(III)** Increasing amounts of kinetochore-microtubule attachments silence the production of the MCC at kinetochores and the APC/C is released. Cdc20 now presents its substrates Cyclin A and Cyclin B (some eukaryotes have other substrates as well, but they are not universally conserved) for ubiquitination and subsequent degradation through recognition of a Dbox motif^42^. Chromosome segregation will now be initiated (anaphase). **B)** Presence/absence matrix of motifs involved in SAC signalling in a selection of Eukaryotes and Metamonads, including *C. membranifera* and *C. frisia*. Colours correspond to the motifs in panel A, light grey indicates motif loss. *N* signifies the number of MadBub homologs that are present in each species. ‘Incomplete’ points to sequences that were found to be incomplete due to gaps in the genome assembly. Question marks indicate the uncertainty in the presence of that particular motif. Although Metamonads have all four MCC components (Mad2, Bub3, MadBub and Cdc20), most homologs do not contain the motifs to elicit a canonical SAC signalling and it is therefore likely that they do not have a SAC response. Exceptions are *C membranifera*, *C. frisia* and *Kipferlia bialata*. They retained the N-terminal KEN-boxes and one ABBA motif, which are involved in the binding of two Cdc20s and a Mad2-interaction motif (MIM) in Mad1 and Cdc20. **C)** Multiple sequence alignments of the motifs from panel A and B. Coloured motif boxes correspond to panel A and B. Multiple sequence alignments were visualized with Jalview^29^, using the Clustal colouring scheme. Asterisks indicate ambiguous motifs in *Carpediemonas membranifera*.

**Supplementary Fig 7.**
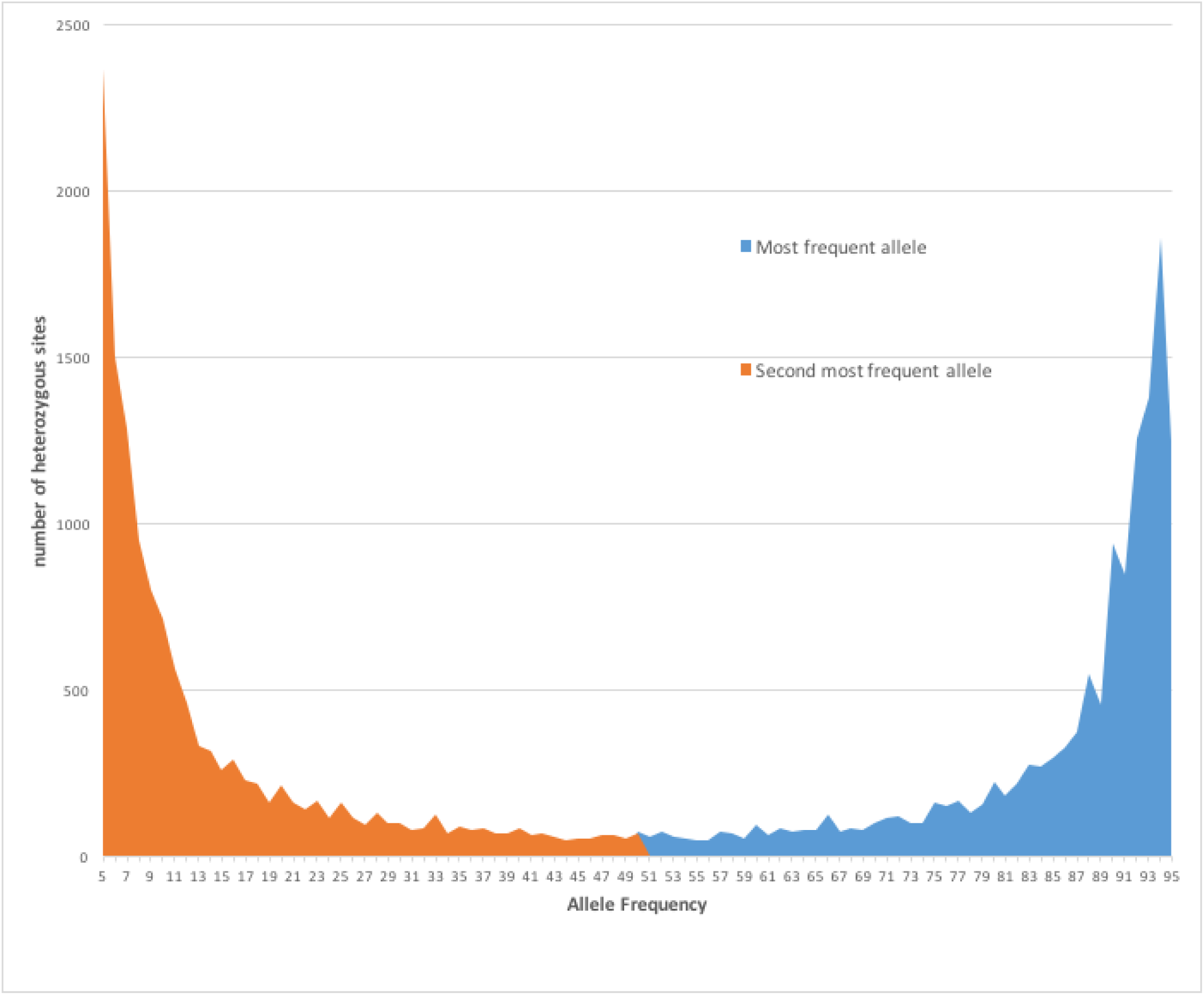
Histogram showing the frequency distribution of single nucleotide variants in the genome of *C. membranifera*. Diagram showing the typical distribution of a haploid genome.

**Supplementary Fig 8.**
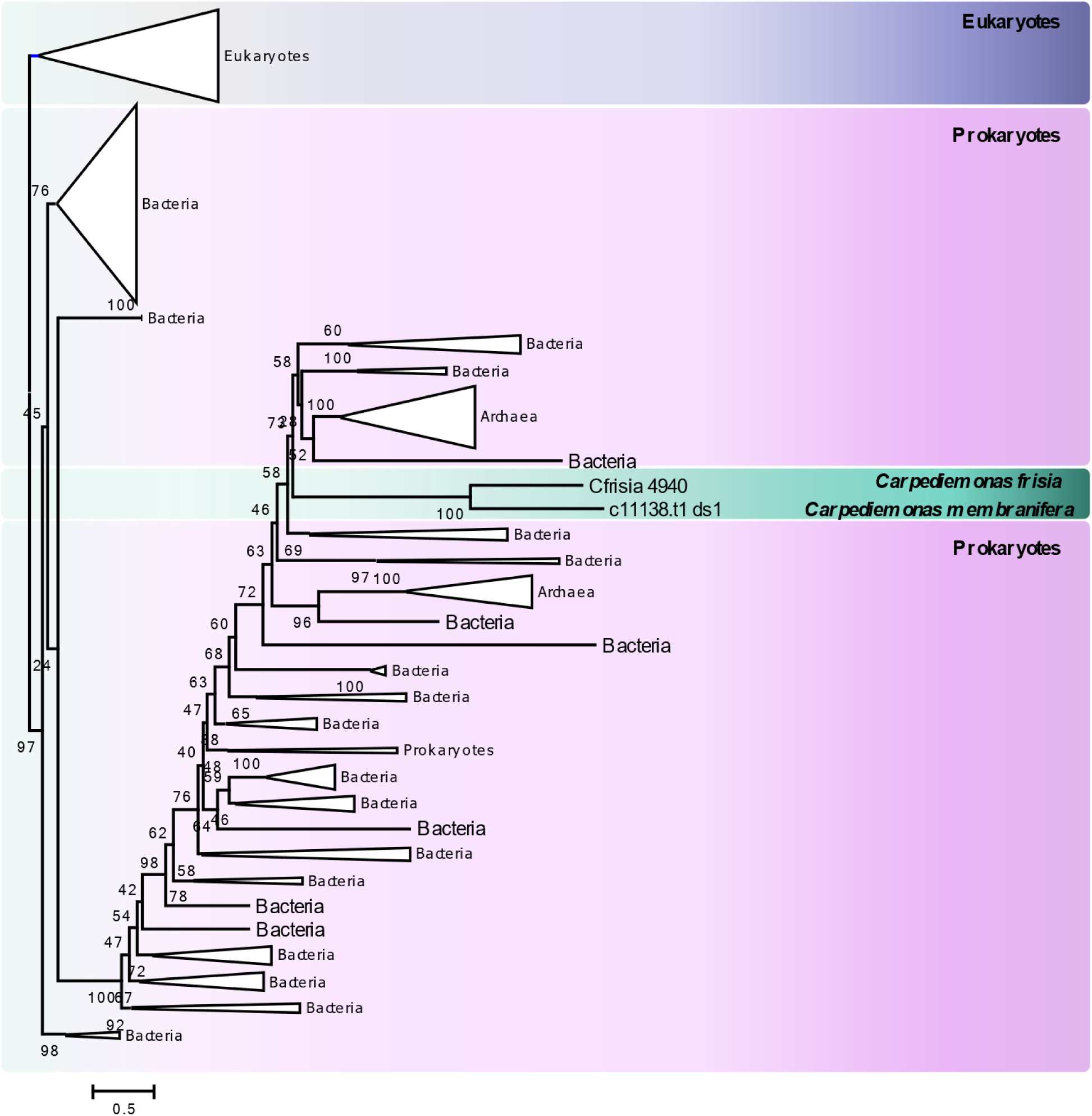
Maximum likelihood reconstruction of Endo IV. The unrooted tree contains eukaryotic and prokaryotic Endo IV sequences, showing *Carpediemonas* sequences emerging within bacterial proteins. The tree was inferred with IQ-TREE under the LG+I+C20 model with 1000 ultrafast bootstraps; alignment length was 276. Scale bar shows the inferred number of amino acid substitutions per site.

**Supplementary Fig 9.**
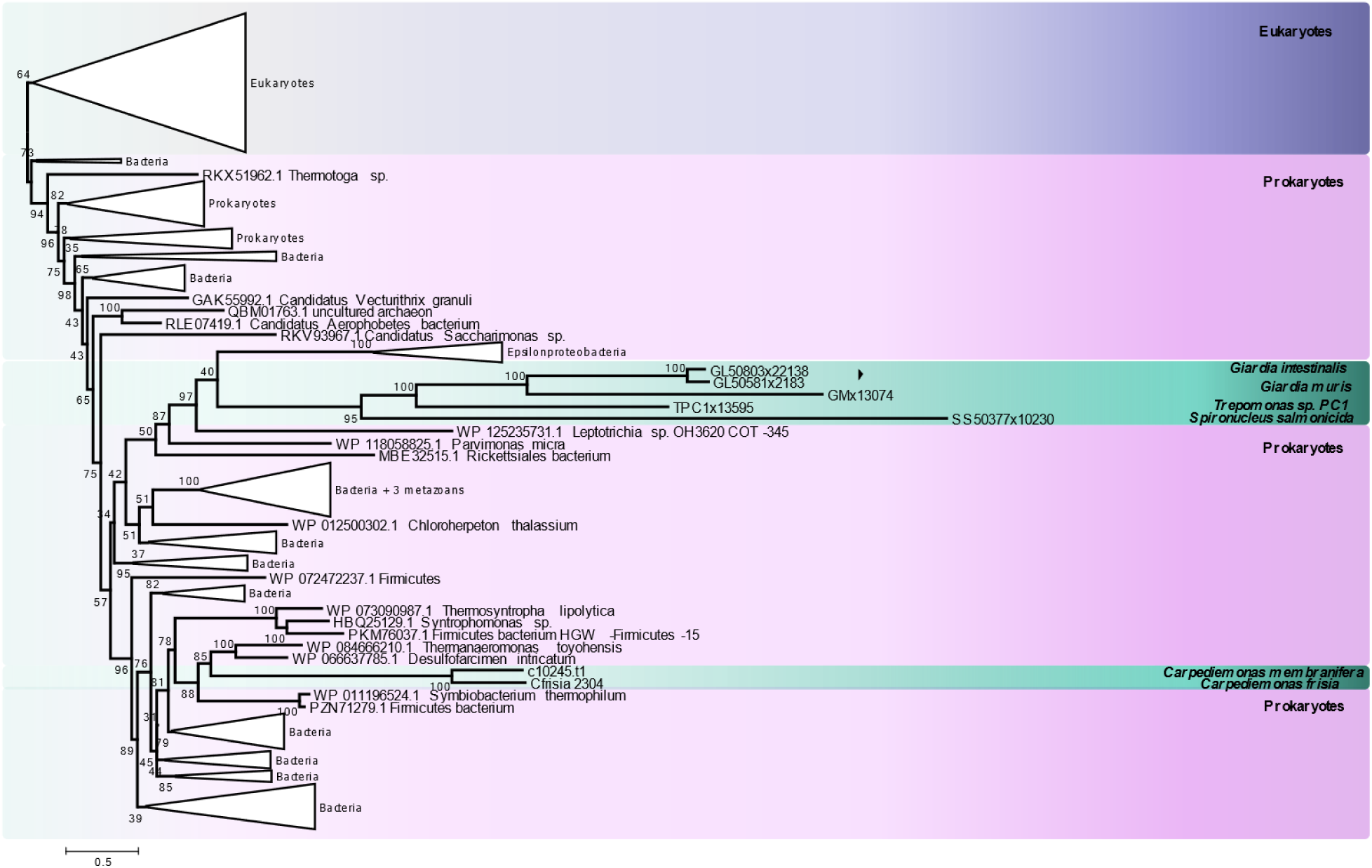
Maximum likelihood reconstruction of RarA. The unrooted tree contains eukaryotic and prokaryotic sequences, showing *Carpediemonas* sequences emerging within bacterial proteins. The tree was inferred with IQ-TREE under the LG+I+C20 model with 1000 ultrafast bootstraps; alignment length was 414. Scale bar shows the inferred number of amino acid substitutions per site.

**Supplementary Fig 10.**
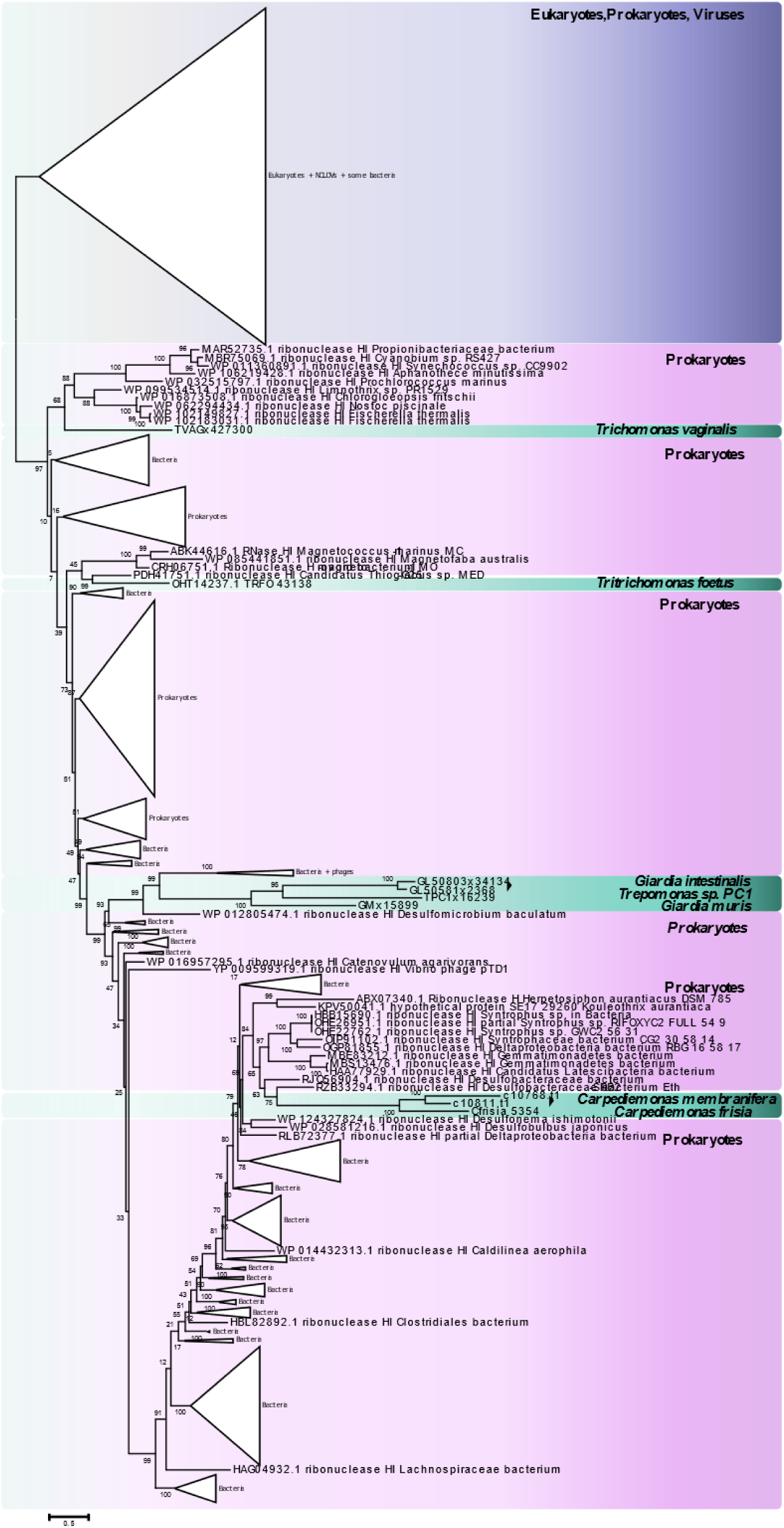
Maximum likelihood reconstruction of RNAse H1. *Carpediemonas* RarA-like proteins emerge within bacterial proteins. Parabasalia and diplomonada proteins highlighting the proteins have been acquired in different events. The tree was inferred with IQ-TREE under the LG+I+G+C20 model with 1000 ultrafast bootstraps; alignment length was 149. Scale bar shows the inferred number of amino acid substitutions per site.

## F. Supplementary tables

Secure download link: http://perun.biochem.dal.ca/downloads/dsalas/Supplementary_Table1.zip

Supplementary Table 1:

**Supplementary Table 1A** BUSCO proteins found in Metamonada based on searches for 245 proteins present in at least one taxon

**Supplementary Table 1B** DNA replication and repair orthologs in 18 diverse eukaryotic genomes

**Supplementary Table 1C** Spindle assembly, kinetochore and APC/C orthologs in 18 diverse eukaryotic genomes

**Supplementary Table 1D** Additional genomes queried during the searches for ORC, Cdc6 and Ndc80 proteins

**Supplementary Table 1E** Lengths of Orc1-6, Cdc6 and Orc1/Cdc6-like proteins and domain architecture comparisons between metamonads and other eukaryotes.

**Supplementary Table 1F** Orc1, Cdc6 and Orc1/Cdc6-likeproteins. Information used in Supplementary Figure 1 panels B and D

